# Evolving antibody response to SARS-CoV-2 antigenic shift from XBB to JN.1

**DOI:** 10.1101/2024.04.19.590276

**Authors:** Fanchong Jian, Jing Wang, Ayijiang Yisimayi, Weiliang Song, Yanli Xu, Xiaosu Chen, Xiao Niu, Sijie Yang, Yuanling Yu, Peng Wang, Haiyan Sun, Lingling Yu, Jing Wang, Yao Wang, Ran An, Wenjing Wang, Miaomiao Ma, Tianhe Xiao, Qingqing Gu, Fei Shao, Youchun Wang, Zhongyang Shen, Ronghua Jin, Yunlong Cao

## Abstract

The continuous evolution of SARS-CoV-2, particularly the emergence of the BA.2.86/JN.1 lineage replacing XBB lineages, necessitates re-evaluation of current vaccine compositions. Here, we provide a comprehensive analysis of the humoral immune response to XBB and JN.1 human exposures, emphasizing the need for JN.1-lineage-based boosters. We demonstrate the antigenic distinctiveness of XBB and JN.1 lineages in SARS-CoV-2-naive individuals but not in those with prior vaccinations or infections, and JN.1 infection elicits superior plasma neutralization titers against its subvariants. We highlight the strong immune evasion and receptor binding capability of KP.3, supporting its foreseeable prevalence. Extensive analysis of the BCR repertoire, isolating ∼2000 RBD-specific monoclonal antibodies (mAbs) with their targeting epitopes characterized by deep mutational scanning (DMS), underscores the systematic superiority of JN.1-elicited memory B cells (MBCs). Notably, Class 1 IGHV3-53/3-66-derived neutralizing antibodies (NAbs) contribute majorly within wildtype (WT)-reactive NAbs against JN.1. However, KP.2 and KP.3 evade a substantial subset of them, even those induced by JN.1, advocating for booster updates to KP.3 for optimized enrichment. JN.1-induced Omicron-specific antibodies also demonstrate high potency across all Omicron lineages. Escape hotspots of these NAbs have mainly been mutated in Omicron RBD, resulting in higher immune barrier to escape, considering the probable recovery of previously escaped NAbs. Additionally, the prevalence of broadly reactive IGHV3-53/3-66- encoding antibodies and MBCs, and their capability of competing with all Omicron-specific NAbs suggests their inhibitory role on the de novo activation of Omicron-specific naive B cells, potentially explaining the heavy immune imprinting in mRNA-vaccinated individuals. These findings delineate the evolving antibody response to Omicron antigenic shift from XBB to JN.1, and highlight the importance of developing JN.1 lineage, especially KP.3-based vaccine boosters, to enhance humoral immunity against current and future SARS-CoV-2 variants.

## Main

Since the emergence of the SARS-CoV-2 BA.2.86 lineage in July 2023, its subvariants, especially JN.1, have continued to circulate and evolve rapidly, outcompeting the previously prevalent XBB subvariants ^1–5^. By June 2024, the JN.1 lineage accounted for over 93% of newly observed sequences (Fig. 1a). Importantly, BA.2.86 and JN.1 have convergently accumulated mutations on the receptor-binding domain (RBD) of the viral spike glycoprotein, including R346S/T, F456L/V, and A475V/S (Extended Data Fig. 1a). A newly detected subvariant, designated as KP.3, even carries an unprecedented Q493E mutation ^6,7^. Most of these sites mutated in JN.1 subvariants are located near the receptor-binding motif (RBM) (Extended Data Fig. 1b). This makes it crucial to investigate their capabilities of evading the current humoral immune barrier established by SARS-CoV-2 infections and vaccines.

**Figure 1.**
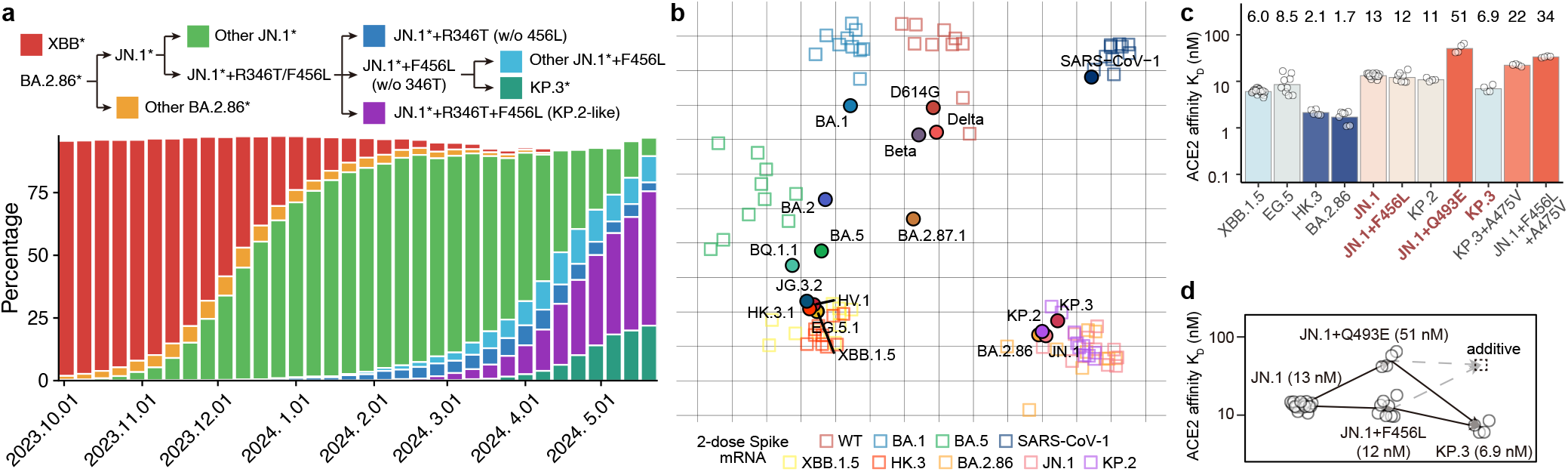
Antigenicity and receptor binding of emerging JN.1 subvariants. **a**, Dynamics of the percentage of XBB and JN.1 lineages in GISAID sequences from Sept 2023 to June 2024. **b**, Antigenic cartography of mouse sera neutralization data with SARS-CoV-2 variant Spike vaccination. Each square indicates a plasma sample and each circle indicates a SARS-CoV-2 variant. **c**, Barplots show the affinities of SARS-CoV-2 variants determined by SPR. Each circle indicates a replicate. Geometric mean K_D_ (nM) values are indicated by height of bars and annotated above each bar. **e**, Schematic for the non-additive ACE2 binding impacts between F456L and Q493E. The dashed gray arrows show the additive result.

Previous studies demonstrated the capability of eliciting JN.1-effective NAbs of XBB-based vaccine boosters ^8–10^. However, considering the distinct antigenicity of JN.1, it is important to investigate whether JN.1 immunization performs substantially better against current and potential future variants ^2,3,11^. Here in this manuscript, we provided a systematic comparison of the humoral immune response between XBB and JN.1 lineages in human infections at both serum and MBC-encoded antibody resolution.

## Results

### Immunogenicity of JN.1 exposure

To evaluate the antigenicity and immunogenicity of the XBB and JN.1 lineages, we first administered a two-dose immunization of variant Spike mRNA in naïve mice (Extended Data Fig. 2a). Our observations revealed a pronounced distinction in antigenicity between the XBB and JN.1 lineages (Fig. 1b and Extended Data Fig. 2b). Notably, within the JN.1 family, KP.3 showed considerable antigenicity difference than JN.1 and KP.2, even when immunizing with KP.2 spike. These distinctions in antigenicity, at least in naive mice, prompts the consideration of changing SARS-CoV-2 vaccine compositions from XBB to JN.1 families.

Future SARS-CoV-2 variant prevalence is a critical guidance for vaccine composition assessment. Human ACE2 (hACE2)-binding affinity of viral RBDs is highly related to viral fitness, and previous studies have highlighted the synergistic impact of RBD L455-F456 mutations on ACE2 receptor binding affinity mediated by Q493 ^12–16^. Given these sites are also convergently mutated in BA.2.86 lineages especially JN.1, we tested the binding affinities of JN.1 subvariant RBD to hACE2 using surface plasmon resonance (SPR) (Extended Data Fig. 1c). Interestingly, F456L (K_D_ = 12 nM) and R346T + F456L (K_D_ = 11 nM) did not largely affect the hACE2-binding affinity of JN.1 (K_D_ = 13 nM), indicating that unlike L455F in HK.3, the dampened ACE2 affinity of JN.1 due to L455S could not be compensated by F456L. Importantly, the Q493E mutation of KP.3 substantially improved the receptor binding affinity on the basis of JN.1 + F456L, exhibiting K_D_ = 6.9 nM (Fig. 1c and Extended Data Fig. 1d). This is not consistent with previous DMS results for ACE2 binding based on BQ.1.1 and XBB.1.5 RBD, with Q493E impairing ACE2 binding, which indicates non-additive epistatic interactions between Q493E and other mutations of KP.3 in comparison with these pre-BA.2.86 variants. Indeed, the Q493E mutation significantly reduces the ACE2 binding affinity in the context of JN.1 RBD (K_D_ = 51 nM), but unexpectedly enhances the affinity when combined with the F456L mutation (Fig. 1d). A475V is a RBM mutation that convergently emerged in multiple lineages, such as JD.1.1, HK.3.14, JN.4, and KP.2.3.1. It decreases ACE2 affinity and escapes NAbs, and therefore, can only be accommodated by variants with high affinity to ACE2 ^3,17,18^. The high affinity of KP.3, achieved through epistasis, may enable the incorporation of A475V for further immune evasion, given the K_D_ is 22 nM for KP.3 + A475V (Fig. 1c). Overall, the extraordinary ACE2-binding affinity may bolster the rapid transmission and prevalence of KP.3, enhancing its potential to acquire additional immune-evasive mutations.

Human serum antibody evasion is the most deciding factor regarding SARS-CoV-2 viral fitness. To analyze the humoral immune evasion capability and immunogenicity of JN.1 lineages, we collected blood samples from 8 cohorts, including individuals infected by XBB* (n=11) or JN.1 (n=4) without known previous exposure to SARS-CoV-2, those who experienced XBB infection after 3 doses of inactivated vaccines, those who experienced sequential infections of BA.5/BF.7 and XBB* (n=14), or BA.5/BF.7 and JN.1 (n=29), and those who received 3-dose inactivated vaccines followed by BA.5/BF.7 breakthrough infection (BTI) and then reinfected by XBB (mainly XBB + S486P), HK.3, or JN.1 (n=54, 18, 29, respectively) (Fig. 2a and Extended Data Fig. 3). Infection strains were inferred based on the sampling time when the corresponding strain was the majority of detected sequences in the region of sample collection (Supplementary Table 1). SARS-CoV-2 Infection was confirmed by either antigen or PCR tests. Plasma samples are isolated and tested for neutralization titers against SARS-CoV-2 variant spike-pseudotyped vesicular stomatitis virus (VSV).

**Figure 2.**
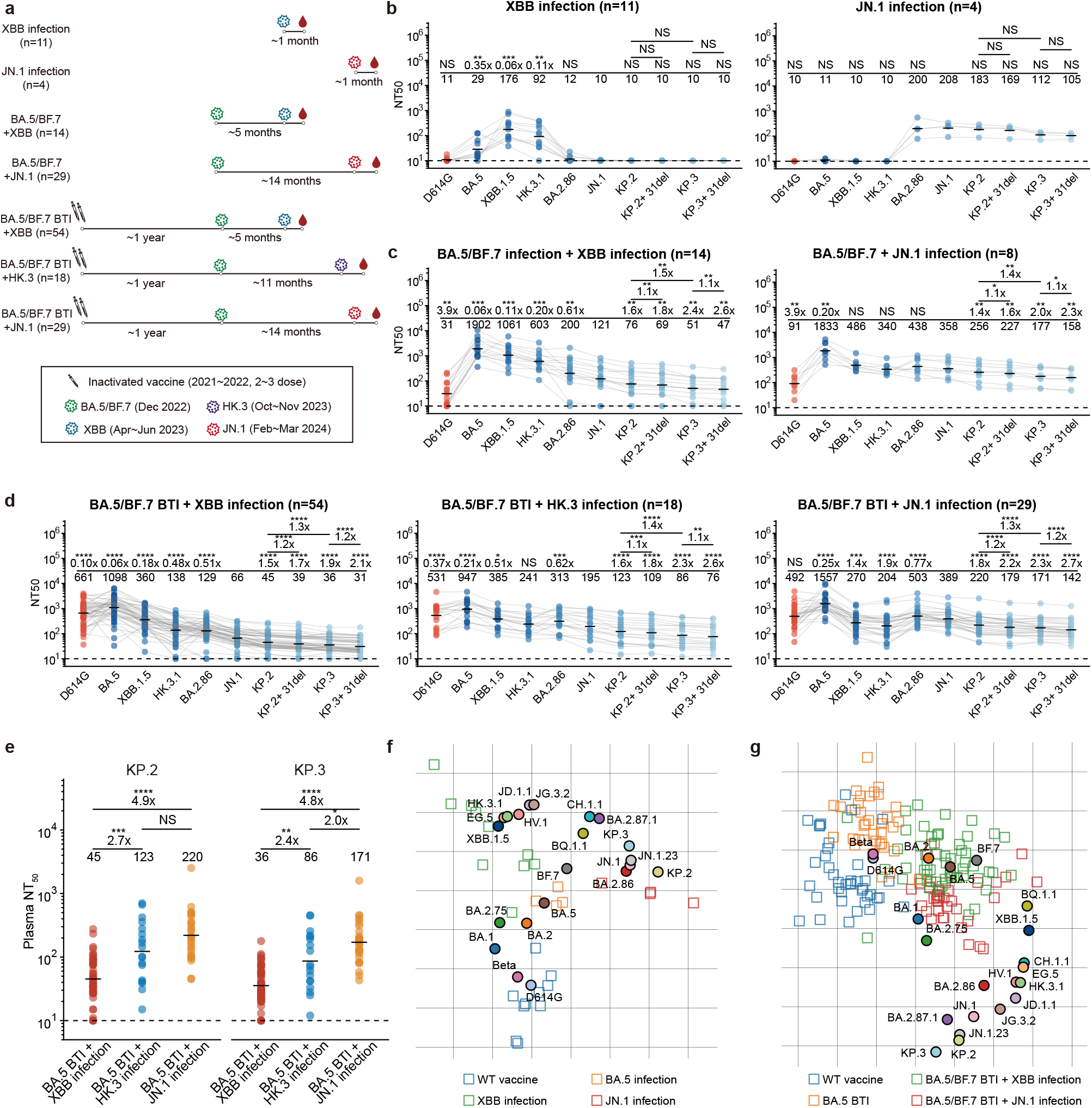
Antigenicity and immunogenicity comparison of XBB and JN.1 lineages in human. **a,** Schematic of the SARS-CoV-2-related immune histories of the seven cohorts involved in this study. **b-d**, 50% neutralization titers (NT_50_) of plasma samples from seven different cohorts against SARS-CoV-2 variant pseudoviruses. Plasma source cohorts and corresponding number of samples are labeled above each panel. Dashed lines indicate the limit of detection (NT_50_ = 10). Numbers of negative samples are labeled below the dashed lines. Geometric mean titers (GMT) values are labeled as black bars and shown above each group of points, with fold-changes and significance compared to JN.1 labeled. Two-tailed Wilcoxon signed-rank tests are used to calculate the p-values. **e**, Comparison of neutralization of plasma samples from three BTI + reinfection cohorts against KP.2 and KP.3. GMT values are labeled as black bars and above the points, with pair-wise fold-changes shown. Two-tailed Wilcoxon rank-sum tests are used to determine the p-values. *p<0.05; **p<0.01; ***p<0.001; ****p<0.0001; NS, not significant. **f-g**, Antigenic cartography performed using human plasma neutralization data of single-exposure cohorts (**f**) or ancestral strain imprinted cohorts (**g**). Each square indicates a plasma sample and each circle indicates a SARS-CoV-2 variant.

Priming with XBB and JN.1 in naïve humans elicited distinct NAbs without observable cross-lineage reactivity, which confirms that XBB and JN.1 and antigenically distinct in both human and mice, indicating that antigenic shift from XBB to JN.1 lineage results in different serotypes (Fig. 2b) ^19,20^. In contrast, a prior BA.5 (or BF.7, omitted hereafter) infection improved the cross-lineage reactivity of antibodies induced by XBB or JN.1 reinfection. This suggests that BA.5/BF.7 priming could induce Omicron cross-reactive NAbs that are effective against both XBB and JN.1 lineages (Fig. 2c).

Notably, in the three BTI with reinfection cohorts, “BA.5 BTI + XBB infection” elicited the lowest NT_50_ against JN.1 lineage variants (Fig. 2d). On average, JN.1 reinfection induced 5.9-fold higher NT_50_ against JN.1, 4.9-fold higher NT_50_ against KP.2, and 4.8-fold higher NT_50_ against KP.3, compared with XBB reinfection (Fig. 2e). The improvement of JN.1 BTI over HK.3 BTI was less significant, possibly due to the shorter interval between two infections in the XBB reinfection cohort, in addition to the immunogenicity drift attributed to the “FLip” mutations (L455F + F456L) of HK.3.

Among all five reinfection cohorts, all of the four tested JN.1 subvariants with RBD mutations, including JN.1 + R346T, JN.1 + F456L, KP.2, and KP.3, exhibited notable immune evasion. KP.3 consistently acted as the strongest escaper, leading to a 1.9 to 2.4-fold reduction in NT_50_ compared to JN.1. A recently emerged deletion on NTD S31, which was convergently detected in multiple independent JN.1 sublineages, including KP.2.3, LB.1, KP.3.1.1, and LF.2, results in slight but significant evasion, which shows the contribution of NTD-targeting NAbs in all six cohorts with repeated SARS-CoV-2 exposure (Fig. 2c-d and Extended Data Fig. 3) ^21^.

Antigenic cartography of our plasma neutralization data visualized the antigenic differences of SARS-CoV-2 variants. The antigenic map from single-exposure cohorts clearly depicted the intrinsic antigenic distances between XBB and JN.1 lineage in human, despite sample size limitations (Fig. 2f). Samples from BTI with reinfection cohorts showed strong ancestral strain imprinting, indicated by the aggregation of points near the D614G strain (Fig. 2g). Nevertheless, the JN.1 BTI cohorts displayed closer distance to current circulating variants, supporting the idea of switching vaccine boosters to JN.1 lineages.

Together, these observations underscore the significant antigenic distinctions between the SARS-CoV-2 XBB and JN.1 lineages, and highlight the notable ACE2 affinity and NAb-escaping capability of emerging JN.1 subvariants, especially KP.3 and KP.3 + S31del (KP.3.1.1). The results provide phenomenological but compelling evidence to shift the focus of vaccine booster strategies from XBB to the JN.1 family.

### JN.1-induced memory B cell repertoire

Then, we aim to dissect the specific molecular constituents responsible for the broad-spectrum neutralization observed in the plasma polyclonal antibodies (pAbs) elicited by infections with the JN.1 lineage, which would enable us to understand how prior vaccination or infection with BA.5 facilitates the development of cross-lineage NAbs following infections with XBB/JN.1. Analyzing the MBC repertoire could also help to predict the response to future variant exposures. Consequently, it is imperative and compelling to deconvolute the roles of antibodies that exhibit diverse cross-reactivities and target multiple epitopes, particularly on the virus RBD, the most immunogenic domain targeted by NAbs.

Therefore, we employed fluorescence-activated cell sorting (FACS) to isolate RBD-specific CD20^+^ CD27^+^ IgM^−^ IgD^−^ B cells using fluorescence activated cell sorting (FACS) from the peripheral blood mononuclear cells (PBMCs) of the human donors previously mentioned. These individuals represented seven distinct SARS-CoV-2-related immune histories: XBB infection, XBB BTI, BA.5 + XBB infection, BA.5 + JN.1 infection, BA.5 BTI + XBB infection, BA.5 BTI + HK.3 infection, and BA.5 BTI + JN.1 infection. We utilized variant RBDs (XBB.1.5, HK.3, or JN.1) corresponding to the last-exposure SARS-CoV-2 strain for each cohort in the sorting (Supplementary Data Fig. 1). Following our previously established methodology, we determined the sequences of the mAb heavy and light chain variable domains using single-cell V(D)J sequencing (scVDJ-seq) and expressed them with human IgG1 Fc backbones ^22–26^. The resultant mAbs were characterized using enzyme-linked immunosorbent assays (ELISA) to assess their binding specificities against the WT and the corresponding Omicron RBDs.

At the resolution of plasma analysis, BA.5 BTI + reinfection consistently produced higher neutralization titers against BA.5 compared to those against D614G, demonstrating the substantial contribution of Omicron-specific NAbs that do not cross-react with the WT (Fig. 2d). This is validated by mAb analyses, in alignment with our earlier discovery that repeated Omicron infections may mitigate the imprinting of inactivated vaccines based on the ancestral strain ^22^. Nevertheless, recent research involving individuals who underwent Omicron reinfection after receiving mRNA vaccines based on the ancestral strain revealed pronounced immune imprinting; as a result, Omicron-specific MBCs were scarcely detectable even after two exposures to Omicron ^27–29^.

The XBB BTI cohort, comprising convalescents who underwent a single Omicron exposure post-vaccination, exhibits the highest proportion (62%) of RBD-specific mAbs that cross-react with the WT. Intriguingly, some vaccine-naïve cohorts, including XBB infection, BA.5 + XBB infection, and BA.5 + JN.1 infection, also generate 40-50% WT-reactive antibodies. The BA.5 + JN.1 infection cohort induces a higher percentage of WT-reactive mAbs compared to the BA.5 BTI + JN.1 infection (Fig. 3a). However, the corresponding plasma samples did not display elevated neutralization titers against the D614G pseudovirus, suggesting an enrichment of cross-reactive mAbs that target non-neutralizing epitopes (Fig. 2c). Remarkably, the BA.5 BTI + HK.3 cohort elicits 74% Omicron-specific mAbs, the highest among the studied groups.

**Figure 3.**
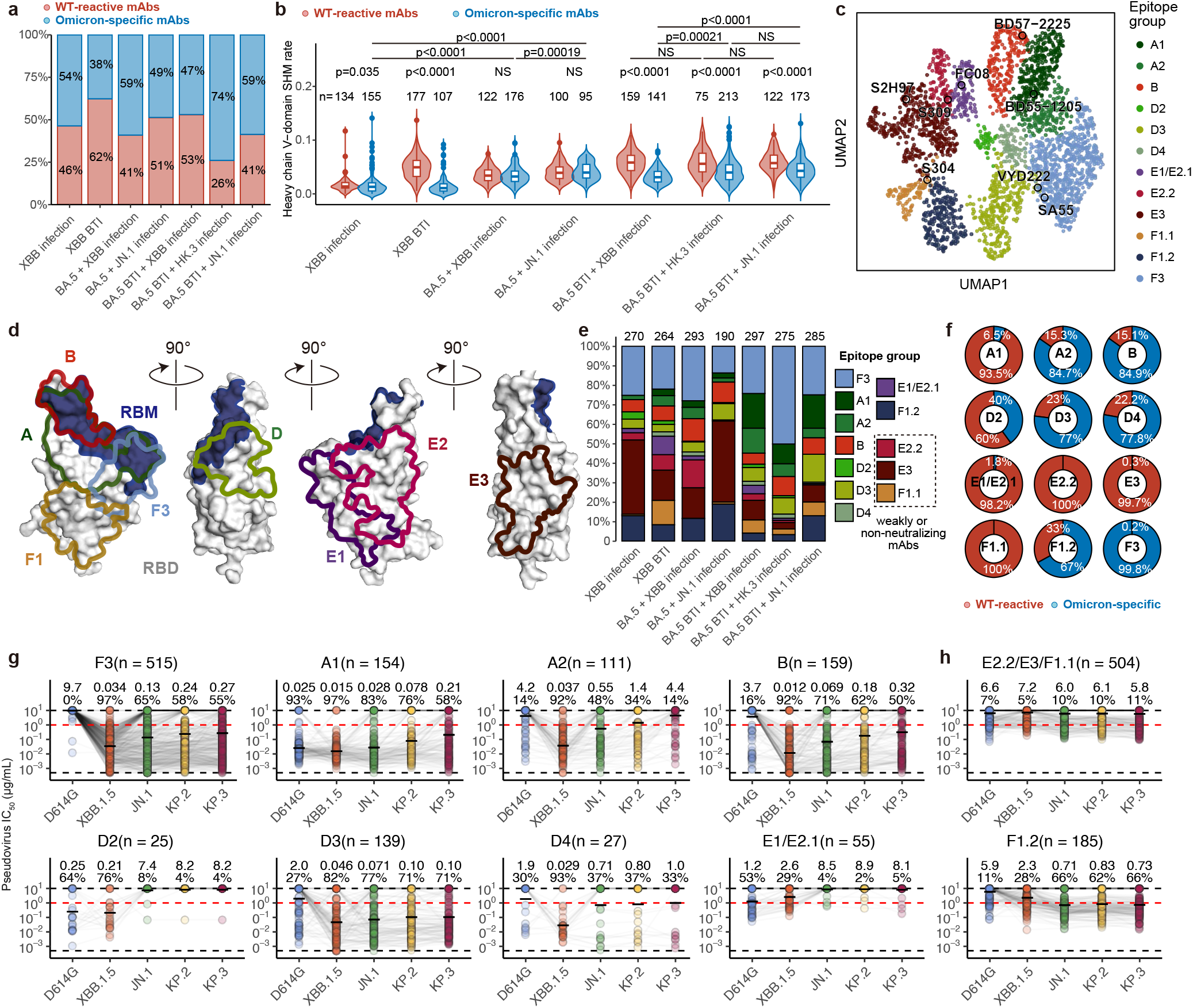
Detailed characterization of XBB/JN.1-elicited mAbs. **a**, Proportion of WT-reactive and Omicron-specific mAbs isolated from different cohorts. Antibody reactivities were determined by ELISA against SARS-CoV-2 WT RBD and XBB.1.5, HK.3, or JN.1 RBD corresponding to the last-exposure variant. **b**, Distribution of heavy chain SHM rate of WT-reactive and Omicron-specific antibodies isolated from different cohorts. Number of mAbs are annotated above each violin plot. Two-tailed Wilcoxon rank-sum tests are used to calculate the p-values. NS, not significant. **c**, Uniform manifold approximation and projection (UMAP) visualization of antibody DMS escape mutation profiles. Well-known mAbs are highlighted by circles with names annotated. **d**, Schematic for the targeting sites of each epitope group on RBD. Epitope groups targeting spatially overlapped epitope are merged. **e**, Percentage of mAbs from each cohort in each epitope group. Number of epitope-characterized mAbs are labeled above the bars. **f**, Percentage of WT-reactive and Omicron-specific mAbs in each epitope group. **g-h**, Neutralization activities in IC_50_ of mAbs in each epitope group against D614G, XBB.1.5, JN.1, KP.2, and KP.3 pseudovirus. Number of mAbs in each group are shown in subtitles. Geometric mean IC_50_ (μg/mL) and percentage of mAbs with IC_50_ < 1 μg/mL are labeled above each group of points.

We observe substantial variations in VDJ gene usage among the mAbs with different reactivities to WT and elicited by different immune histories. In the BA.5 BTI + reinfection cohorts, there is a prominent usage of IGHV3-53/3-66 in WT-reactive mAbs, which are recognized for being part of the public immune response and predominantly Class 1 NAbs targeting the RBM. However, this type of mAbs is scarcely seen in cohorts without vaccination, where there is a higher utilization of IGHV5-51 and IGHV4-39 (Extended Data Fig. 4a). Regarding Omicron-specific mAbs, IGHV2-5 is prevalent across all cohorts, yet interestingly, it is not dominant among JN.1-infected convalescents, who show a higher proportion of mAbs derived from IGHV5-51 (Extended Data Fig. 4b). Notably, IGHV5-51 is also extensively used in WT-reactive antibodies, underscoring its significance, particularly in the context of JN.1 infections.

As expected, the rates of somatic hypermutation (SHM) in both the heavy and light chains of mAbs are closely associated with the number of antigen exposures. Specifically, WT-reactive mAbs exhibit more SHMs than Omicron-specific mAbs in vaccinated individuals, but not in unvaccinated ones. The cohorts BA.5 BTI + HK.3/JN.1 generate Omicron-specific mAbs with higher SHM rates compared to BA.5 BTI + XBB infection, likely due to the longer interval between two Omicron exposures in the former groups, allowing for further maturation of Omicron-specific B cells initiated by BA.5 infections (Fig. 3b and Extended Data Fig. 4c).

We further evaluated the neutralization capabilities of the derived mAbs against variant pseudoviruses. Generally, Omicron-specific mAbs demonstrated superior neutralization activities compared to WT-reactive mAbs against variants JN.1, KP.2, and KP.3. mAbs induced by XBB infection and XBB BTI displayed an exceedingly low percentage of potent NAbs, consistent with their low plasma neutralization titers (Fig. 2a and Extended Data Fig. 3a). Notably, BA.5 + JN.1 and BA.5 BTI + JN.1 infections elicited 30% and 60% JN.1-effective WT-reactive NAbs, respectively, while the proportion of effective Omicron-specific mAbs exceeded 90% in both cohorts, surpassing those observed in XBB/HK.3 reinfections (Extended Data Fig. 4d). These findings further substantiate the potential benefits of developing vaccine boosters based on the JN.1 lineages.

### Epitope mapping of JN.1-induced mAbs

Despite the promising neutralization activities of JN.1-elicited mAbs, antibodies targeting various epitopes may be evaded by diverse RBD mutations, suggesting their potential vulnerability to future viral antigenic drift. To further elucidate the molecular components and mechanisms underlying the broadly neutralizing antibodies (bnAbs) induced by XBB/JN.1 infections at the resolution of single amino acid epitopes, we conducted high-throughput yeast-display-based DMS assays to analyze the escape mutation profiles of the isolated mAbs. Specifically, we constructed mutant libraries based on the XBB.1.5 and JN.1 RBDs. We initially assessed the expression levels of these mutants on the yeast surface using FACS followed by sequencing (Sort-seq) (Extended Data Fig. 5a-d) ^6,7,30^. Interestingly, the expression of the JN.1 RBD appeared to be more tolerant to mutations compared to the BA.2 RBD, yet less tolerant than the XBB.1.5 RBD (Extended Data Fig. 5e). Following our established protocol, we conducted DMS on the mAb binding capabilities using magnetic beads-based sorting in a high-throughput manner to identify the escape mutations for each mAb, thereby mapping their targeting epitopes ^22^. We successfully assayed the escape mutation profiles of a total of 2,688 mAbs, based on at least one of the two RBD variants, including 1,874 isolated from XBB/JN.1 infection cohorts involved in this study, and 814 mAbs previously identified by other groups or ourselves for comparison (Extended Data Fig. 6a) ^22,24,31^. We utilized a graph-based unsupervised clustering algorithm to identify 22 mAb clusters, and the corresponding epitope groups for each cluster were annotated based on our previous definitions (Fig. 3c and Extended Data Fig. 6b) ^22,24^. In brief, epitope groups A1/A2 (Class 1^32,33^), B (Class 1/2, similar to COV2-2196^34^ and REGN10933^35^), D2/D3/D4 (similar to REGN10987^35^ and LY-CoV1404^36^), and F3 (Class 1/4, similar to SA55^37^ and ADG-2/VYD222^38^) generally compete with the receptor ACE2, and have a higher potential to effectively neutralize the virus. Conversely, groups E1/E2 (Class 3, exemplified by S309), E3 (also referred to as “Class 5”, exemplified by S2H97^39^), and F1 (Class 4, exemplified by S304) are less likely to compete with ACE2 and exhibit potent neutralization (Fig. 3d and Extended Data Fig. 6c-d). Notably, we discovered a novel subgroup of F1, designated as F1.2, which targets an epitope adjacent to the traditional F1.1 but slightly closer to the RBM (Extended Data Fig. 6e).

This comprehensive epitope mapping allows for the examination of the epitope distribution of mAbs isolated from human donors with varying immune histories (Fig. 3e). We observe that the proportion of A1 mAbs correlates with the number of SARS-CoV-2 exposures, reaching highest levels in cohorts that experienced BTI followed by reinfection. However, these antibodies are notably absent in cases of XBB infection alone, underscoring the importance of initial exposure to earlier variants for the development of such mAbs. By differentiating F1.2 from F1.1, we deduce that WT-based vaccination is essential for eliciting traditional F1.1 non-neutralizing antibodies. In contrast, immunization solely with Omicron induces F1.2 mAbs only, which may be due to the immunogenicity shift caused by Omicron mutations at RBD positions 371-376, rendering F1.2 immunodominant and masking the F1.1 epitope. We also note that JN.1 infections do not elicit E1/E2 mAbs, which could be attributed to the N354 glycosylation resulting from the K356T mutation in the BA.2.86 lineage. Notably, HK.3 reinfection induces a substantial proportion of F3 mAbs (Fig. 3e). Among the epitope groups, A1, D2, E1/E2/E3, and F1.1 are predominantly cross-reactive to WT; whereas A2, B, D3/D4, F1.2, and F3 primarily consist of Omicron-specific mAbs (Fig. 3f). Groups F3, A1, B, and D3 encompass potential bnAbs against JN.1 subvariants, whereas A2, D2, D4, and E1/E2.1 are largely escaped (Fig. 3g). E2.2/E3/F1.1 typically represent broadly reactive non-neutralizing antibodies. However, the novel F1.2 mAbs, which exhibited weak neutralization against SARS-CoV-2 variants prior to BA.2.86, demonstrate an unprecedentedly enhanced potency against JN.1 lineages (Fig. 3h). In addition, we find groups E1/E2.1, E2.2, F1.1, and F1.2 show a significant preference for light chain V genes, enriching for IGLV1-40, IGLV3-21, IGKV1-39, and IGLV6-57, respectively. Additionally, E1/E2.1 and F1.1 tend to utilize IGHV1-69 and IGHV3-13/3-30 heavy chains, respectively, to pair with their corresponding light chains (Extended Data Fig. 6f).

### Class 1 mAbs dominate WT-reactive bnAbs

Given the potential scarcity of Omicron-specific NAbs within the mRNA-vaccinated population, to explore the real-world JN.1 lineage evolution and its interaction with population-level humoral immunity, and to determine whether JN.1 infection could efficiently induce WT-reactive NAbs that neutralize JN.1 subvariants, we then focus on the properties of WT-reactive mAbs elicited by three BA.5 BTI + reinfection cohorts. Consistent with the plasma neutralization and overall mAb neutralization analyses shown above, WT-reactive mAbs from HK.3 and JN.1 infections were significantly more effective than those from XBB infection against JN.1, KP.2, and KP.3 (Fig. 4a). We then calculated the “effectiveness scores” for each epitope group from each source cohort, defined as the number of mAbs in each epitope group weighted by their IC_50_ values against a specific variant. This metric helped us discern the contribution of each epitope group to neutralization (Fig. 4b). Notably, epitope group A1 consistently made a major contribution to the effectiveness against not only JN.1 but also KP.2 and KP.3, which accumulate multiple mutations on the A1 epitope or even its escape hotspots, including L455S, F456L, and Q493E (Fig. 4b-c and Extended Data Fig. 6d). WT-reactive mAbs from epitope groups B and D3 also partially contribute to the broad neutralization against JN.1 lineages in the HK.3 and JN.1 reinfection cohorts, despite these cohorts having a high proportion of Omicron-specific mAbs due to the T478K and Q498R/N440K mutations on their epitopes that have been convergently found in all Omicron lineages, respectively (Fig. 3f and 4b-c).

**Figure 4.**
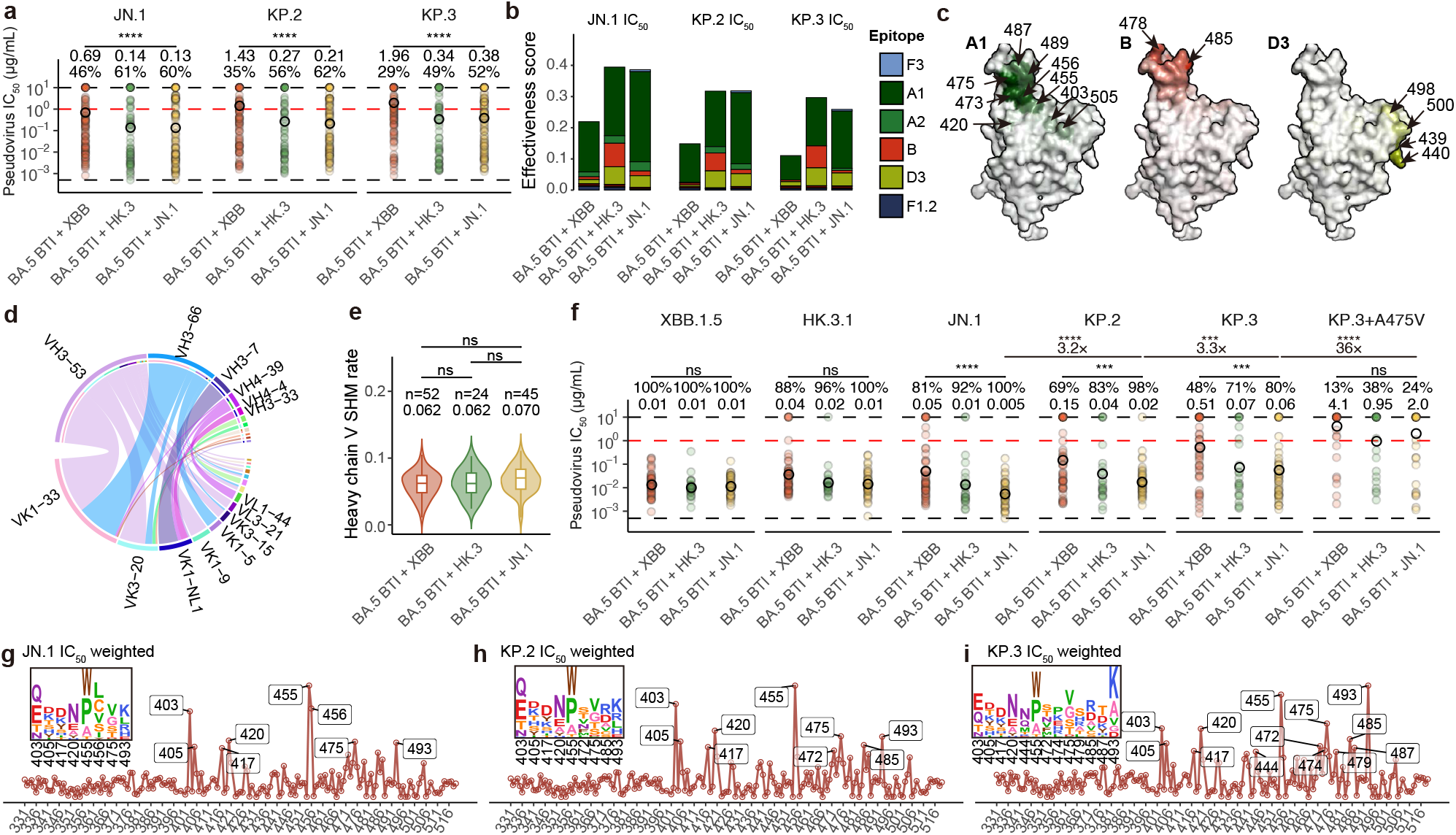
Class 1 dominates WT-reactive bnAbs. **a**, Neutralization activities of WT-reactive mAbs isolated from three BTI + reinfection cohorts against JN.1, KP.2, and KP.3. Geometric mean IC_50_ (μg/mL) and percentage of mAbs with IC_50_ < 1 μg/mL are labeled above each group of points. **b**, Stacked bar charts show the effectiveness scores of WT-reactive mAbs in each epitope group weighted by IC_50_ against JN.1, KP.2, and KP.3. **c**, Average DMS site escape scores of mAbs in epitope groups A1, B, and D3. Hotspot residues are indicated by arrows. **d**, Chord diagram shows the heavy-light chain V gene pairing of mAbs in epitope group A1. The names of corresponding germline genes are annotated next to the strips. **e**, Comparison of heavy chain SHM rate of A1 mAbs elicited by BA.5 BTI + XBB, HK.3, and JN.1 reinfection cohorts. **f**, Neutralization activities of WT-reactive mAbs in epitope group A1 isolated from three BTI + reinfection cohorts. Geometric mean IC_50_ (μg/mL) and percentage of mAbs with IC_50_ < 1 μg/mL (red dashed lines) are labeled above each group of points. Black dash lines indicate limits of detection (0.005 and 10 μg/mL). **g-i**, Calculation of immune pressure on each RBD site and mutation based on the average of WT-reactive antibody escape mutation profiles weighted by JN.1 (**g**), KP.2 (**h**), KP.3 (**i**), and DMS for RBD expression and ACE2 binding. Hotspot residues are labeled and shown in logo plots. Two-tailed Wilcoxon rank-sum tests or signed-rank tests (first row, for paired samples) are used to determine the p-values. ***p < 0.001; ****p < 0.0001. ns, not significant.

In line with previous findings, group A1 or Class 1 mAbs are predominantly derived from the IGHV3-53 or IGHV3-66 germline, which is well-known as the public or convergent humoral immune response to SARS-CoV-2 ^26,33,40,41^. These mAbs tend to pair with the IGKV1-33 light chain; however, a special subset of A1 mAbs from the BA.5 BTI + XBB reinfection cohort utilizes IGHV3-7 with IGKV1-NL1. The WT-reactive A1 mAbs from the three BTI + reinfection cohorts exhibit similar heavy chain SHM rates and similar neutralizing activities against XBB.1.5 and HK.3.1 (Fig. 4e-f). Nonetheless, those elicited by HK.3 and JN.1 demonstrate significantly enhanced neutralization against JN.1 subvariants. Notably, KP.2 and KP.3 evade (IC_50_ > 1 μg/mL) 31% and 52% of the mAbs elicited by XBB reinfection, respectively, but only 2% and 20% of the mAbs elicited by JN.1 reinfection. Compared to those elicited by XBB, JN.1-elicited A1 mAbs exhibit, on average, 7-10-fold higher neutralizing activity against JN.1, KP.2, and KP.3, with an even greater difference than that observed in plasma neutralization titers (Fig. 4f and Extended Data Fig. 7a). Thus, in the context of WT-cross-reactive antibodies, JN.1 infection not only elicits higher neutralization against current JN.1-derived strains but also better enriches MBCs that encode effective Class 1 or epitope group A1 receptor-mimicking antibodies. However, JN.1-elicited WT-reactive A1 NAbs exhibit a 3.2-fold and 10-fold reduction in reactivities against KP.2 and KP.3, respectively. Most strikingly, only 24% retain their neutralization against KP.3 + A475V (Fig. 4f). This susceptibility raises concerns regarding the effectiveness of JN.1 boosters in counteracting ongoing viral evolution, and indicates the need for vaccines derived from the KP.3 lineage for robust protection against both current variants and future antigenic drifts.

We observed that the A1 NAbs broadly against the six tested strains do not exhibit significantly higher SHM rates and do not show a significant preference in the usage of heavy chain and light chain germlines compared to the A1 NAbs that are evaded by at least one variant (Extended Data Fig. 7b-c) ^42^. The escaped A1 mAbs exhibit higher DMS escape scores than the broadly neutralizing A1 mAbs on the mutations of interest, such as 456L and 475V on both antigen basis, but 455S and 493E only on the XBB.1.5 basis. JN.1 binding filtered out the A1 mAbs that are evaded by 455S, and Q493E can be largely influenced by its epistasis with F456L, as mentioned in the ACE2-binding characterization above (Extended Data Fig. 7d-e). Previous structural analyses indicated that IGHV3-53/3-66 mAbs primarily utilize their CDR-H1 and part of CDR-H3 to interact with RBD residue A475; however, we did not observe notable differences in CDR-H1 patterns (Extended Data Fig. 7f) ^4,40^. Therefore, we hypothesize that these A1 bnAbs rely on a distinctive core CDR-H3 and highly matured light chain for broad neutralization, as suggested by the preferences in IGHD gene usage (Extended Data Fig. 7g-h). However, due to the limited number of mAbs, we cannot specifically identify the detailed motif responsible for broad neutralization.

Recent growth advantages of JN.1 subvariants with mutations on A1 epitope indicate the remarkable abundance of such NAbs within the global population. This is also revealed by assessing the average immune pressure by aggregating mAb DMS profiles in a neutralization-weighted, codon-aware manner. We employ WT-reactive NAbs elicited by BTI + reinfection, which should reflect the consensus scenario in the real world. Despite the accumulation of escape mutations on the A1 epitope and the verified significant evasion, the retained A1 bnAbs still exert pressure on residues within its epitope hotspots, such as 403, 420, 455, 475, and 493 (Fig 4g-i). Unsurprisingly, the F456L mutation in KP.2 and KP.3 eliminates the L456 hotspot observed in JN.1 weighting; however, the score on residue E493 is even more pronounced in KP.3 weighting, as this mutation enables four new one-nucleotide-accessible amino acid mutations at this site, including Ala, Asp, Gly, and Val. Due to the absence of ACE2 binding DMS data on the JN.1 basis, we utilized XBB.1.5-based results to filter out ACE2-dampening mutations in our calculations, which may introduce artifacts when strong epistasis is present.

In summary, within the WT-reactive NAbs, epitope group A1 remains the most pronounced against JN.1 subvariants, despite multiple evasive mutations on its epitope during recent viral evolution. Therefore, the development of boosters based on JN.1, or even JN.1 + F456L, KP.2, or KP.3, should be considered to better elicit bnAbs and enrich for effective MBCs that can resist potential future immune escape mutations, particularly in individuals receiving mRNA vaccines, whose immune responses predominantly elicit WT-reactive antibodies.

### Potential of Omicron-specific NAbs

Unlike mRNA vaccination, immune imprinting caused by inactivated vaccines appears to be mitigated by Omicron reinfection, which elicits a substantial amount of Omicron-specific antibody. As global vaccination strategies shift away from WT components and update to the latest variants, such mAbs may become the primary contributors to immune pressure in the future. Notably, JN.1 infection also induces Omicron-specific NAbs with significantly enhanced neutralization breadth against the JN.1 lineage compared to XBB or HK.3 infections (Fig. 5a). Epitope group F3 stands out as the most remarkable for broad neutralization, while A2, B, D3, and F1.2 also make minor contributions (Fig. 5b). A2 NAbs are likely to be evaded due to their highly overlapping epitope with group A1. Interestingly, groups B and D3 include both WT-reactive and non-reactive bnAbs (Fig. 4b and 5b). Unsurprisingly, BTI cohorts elicit more WT-reactive B/D3 mAbs than unvaccinated cohorts, and these cross-reactive B/D3 mAbs exhibit a higher SHM rate than Omicron-specific ones (Extended Data Fig. 8a-b). These findings suggest that at least a subset of these cross B/D3 mAbs may originate from the recall of pre-Omicron memory; however, it is also possible that the higher SHM rate is merely the result of selection for random cross-reactivity in Omicron-specific antibodies. Despite their cross-reactivity to WT, these antibodies demonstrate much higher neutralization activities against BA.5 compared to D614G, indicating potential Omicron-adaptive maturation (Extended Data Fig. 8c). WT-reactive and Omicron-specific B mAbs are derived from largely different heavy and light chain genes; however, D3 mAbs predominantly utilize IGHV5-51 regardless of their cross-reactivity (Extended Data Fig. 8d-e). Specific group B mAbs exhibit higher DMS escape scores on residues 478 and 486, and D3 higher on 440, which are mutated sites in Omicron lineages (Extended Data Fig. 8f-g). Many Omicron-specific B NAbs (but not D3) do not neutralize BA.1 and BA.2, which do not harbor the F486V/S/P mutations found in post-BA.5 variants, due to their vulnerability to 486 mutations (Extended Data Fig. 8h).

**Figure 5.**
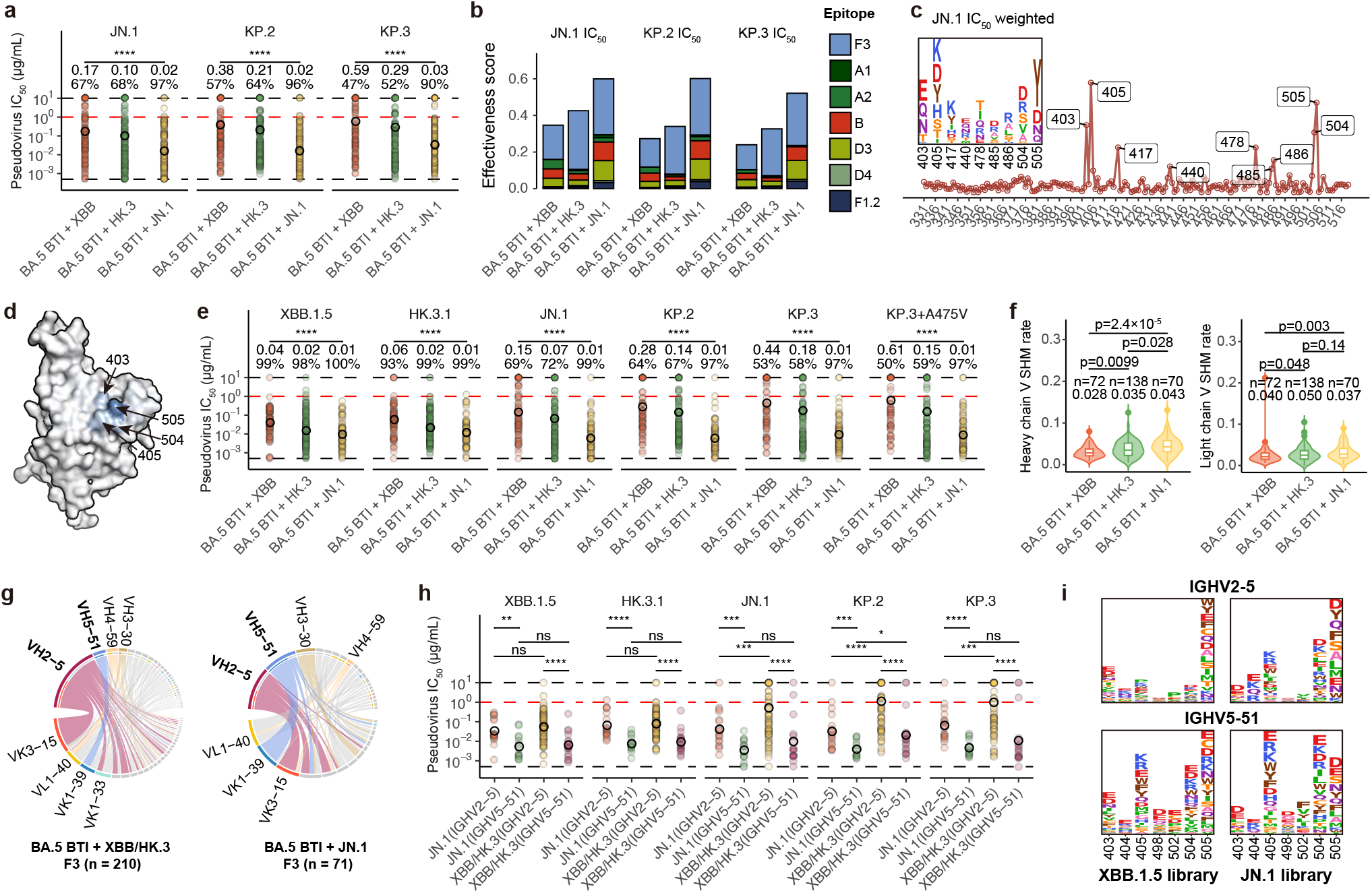
Broad neutralization of Omicron-specific Nabs. a, Neutralization activities of Omicron-specific mAbs isolated from three BTI + reinfection cohorts against JN.1, KP.2, and KP.3. Geometric mean IC_50_ (μg/mL) and percentage of mAbs with IC_50_ < 1 μg/mL are labeled above each group of points. **b**, Stacked bar charts show the effectiveness scores of Omicron-specific mAbs each epitope group weighted by IC_50_ against JN.1, KP.2, and KP.3. **c**, Calculation of immune pressure on each RBD site and mutation based on the average of Omicron-specific antibody escape mutation profiles weighted by JN.1, and DMS for RBD expression and ACE2 binding. Hotspot residues are labeled and shown in logo plots. **d**, Average DMS site escape scores of mAbs in epitope group F3. Hotspot residues are indicated by arrows. **e**, Neutralization activities of Omicron-specific mAbs in group F3 isolated from BTI + reinfection cohorts. Geometric mean IC_50_ (μg/mL) and percentage of mAbs with IC_50_ < 1 μg/mL (red dashed lines) are labeled above each group of points. Black dash lines indicate limits of detection (0.005 and 10 μg/mL). **f**, Comparison of SHM rates of F3 mAbs elicited by BA.5 BTI + XBB, HK.3, and JN.1 reinfection cohorts. **g**, Chord diagram shows the heavy-light chain V gene pairing of mAbs isolated from BA.5 BTI + XBB/HK.3 or JN.1 in epitope group F3. **h**, Neutralization activities of Omicron-specific mAbs in group F3 isolated from BA.5 BTI + XBB/HK.3 or JN.1 cohort encoded by IGHV2-5 or IGHV5-51. **i**, Average DMS escape mutation scores of F3 mAbs encoded by IGHV2-5 or IGHV5-51.

Despite the abundance of potent Omicron-specific NAbs in individuals who have experienced reinfection following prior inactivated vaccinations, we observe minimal evidence of mutations that enable escape from these NAbs. The lack of escape mutations against such NAbs is particularly notable in China, where the majority of the population has received inactivated vaccines combined with BA.5/BF.7 BTI, or even experienced more reinfections, suggesting weak selective pressure or inherent evolutionary constraints that limit the emergence of escape mutations. Through the aggregation of DMS profiles of Omicron-specific NAbs, we have identified that all escape hotspots, except for G504, are located on residues of the RBD that have mutated in Omicron variant (Fig. 5c). Given the potential for neutralization recovery of previously escaped NAbs, these mutated sites may have a reduced likelihood for further mutation. Notably, mutations at G504 have been recently reported to enhance serum neutralization, likely due to their regulatory impact on the up-down dynamics of the Spike glycoprotein ^43^. As anticipated, the four most prominent hotspots, encompassing residues 403, 405, 504, and 505, are all targeted by epitope F3 (Fig. 5d). Also, NAbs induced by JN.1 exhibit a remarkable breadth of neutralization against all tested JN.1 subvariants, outperforming those induced by XBB and HK.3 (Fig. 5e). This superiority is not surprising, as no additional mutations have occurred on their escape hotspots following the R403K mutation in BA.2.86. The SHM rates observed in HK.3 and JN.1-induced F3 mAbs are higher than those induced by XBB, inconsistent with the neutralization results which show that XBB and HK.3 exhibit similar neutralization capabilities. This discrepancy suggests that maturation is not the predominant factor determining the broad neutralization efficacy of F3 (Fig. 5f).

Instead, F3 mAbs display intriguing patterns in the utilization of heavy and light chain V genes. F3 antibodies elicited by a single Omicron exposure, such as XBB infection and XBB BTI cohorts, are almost exclusively derived from the IGHV2-5 and IGKV3-15 pairing (Extended Data Fig. 9a). However, these NAbs exhibit weak neutralization against JN.1 lineages (Extended Data Fig. 9b). In contrast, repeated Omicron infections diversify the germline utilization of F3 mAbs, and generate F3 mAbs of comparable breadth regardless of vaccination (Fig. 5g and Extended Data Fig. 9c-d). Notably, JN.1 infection emphasizes the usage of IGHV5-51, particularly when paired with IGKV1-39. We demonstrate that, regardless of the source cohort, IGHV5-51 F3 antibodies are significantly more effective against JN.1 lineages than IGHV2-5-derived ones (Fig. 5h). However, we did not observe lower DMS scores on residue 403, which is intuitive given its presence in all BA.2.86 subvariants. Conversely, these IGHV5-51 F3 broad bnAbs exhibit higher escape scores on residues 405 and 504 (Fig. 5i). IGHV5-51 appears to be a noteworthy germline heavy chain V gene in the context of the antigenicity and immunogenicity of the JN.1 lineage. Specifically, IGHV5-51 encompasses three major epitope groups, E3, D3, and F3, with distinct patterns of light chain usage. E3 and F3 favor IGLV1-44 and IGKV1-39, respectively, while IGHV5-51 D3 mAbs utilize a wide range of light chain V genes (Extended Data Fig. 9e). The SHM rates of these groups do not show significant differences, and their neutralization capabilities closely align with the properties of their respective epitope groups (Extended Data Fig. 9f-g). A well-known mAb isolated from a SARS convalescent, CR3022, and two recently reported bnAbs isolated from ancestral strain convalescents, are also encoded by IGHV5-51 germline, but paired with light chain germlines not highlighted here^44,45^.

These findings underscore the superior efficacy of JN.1-elicited Omicron-specific NAbs and emphasize the potency of these NAbs, especially the IGHV5-51-encoding F3 NAbs, against all Omicron variants, which should be considered as potential targets for the development of vaccines.

### Clash of Class 1 and Omi-specific NAbs

Recent research has highlighted an exceptionally strong immune imprinting in individuals vaccinated with mRNA vaccines, as they fail to mount an Omicron-specific antibody response even following multiple Omicron exposures ^27–29^. However, this phenomenon cannot be observed in individual who received inactivated vaccines, or in mRNA-vaccinated mice ^27^. Upon the comprehensive characterization of Omicron-specific antibodies, we surprisingly discovered that all Omicron-specific neutralizing epitopes on RBD compete with the A1 mAbs, which are well-known for the convergent usage of IGHV3-53/3-66 germline (Fig. 6a-b). This competition was confirmed by SPR-based competition assays (Fig. 6c). Given these results, we hypothesize that the presence of the IGHV3-53/3-66 convergent response is pivotal for this robust imprinting ^46,47^.

**Figure 6.**
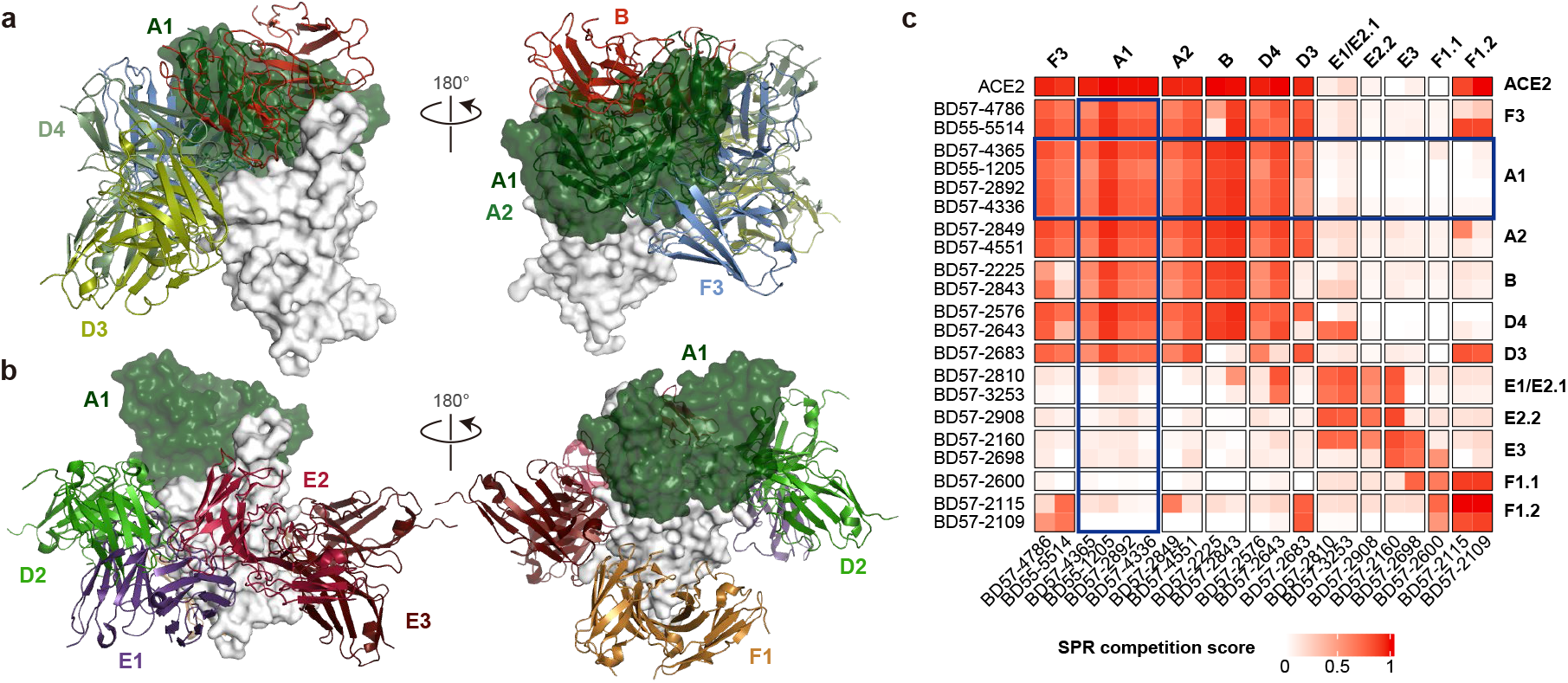
Competition between Class 1 and Omicron-specific NAbs. **a**, Superimposed structural models of representative antibodies in epitope group A1 and Omicron-specific neutralizing epitope groups. **b**, Superimposed structural models of representative antibodies in epitope group A1 and WT-reactive epitope groups. **c**, Heatmap for pair-wised SPR competition scores of representative mAbs in various epitope groups on XBB.1.5 RBD. Results related to epitope group A1 are highlighted by blue rectangles.

In essence, inactivated vaccines induce a weaker convergent response compared to mRNA vaccines. The individuals studied in our research experienced the “zero COVID” period in China during 2021-2022, leading to significant antibody waning. As a result, the concentration of Omicron-effective IGHV3-53/3-66 NAbs and corresponding MBCs may have been insufficient to effectively mask the antigen upon initial exposure to Omicron. This scenario would allow Omicron-specific naïve B cells to be activated and promoted to mature. These activated B cells could then be recalled by a subsequent Omicron exposure, leading to the generation of extensive Omicron-specific MBCs and antibodies. In contrast, the strong convergent responses in mRNA-vaccinated individuals may efficiently mask all Omicron-specific epitopes during the first Omicron encounter ^48^. Their MBCs encoding effective IGHV3-53/3-66 public antibodies would be repeatedly activated with each Omicron exposure, demonstrating remarkable immune imprinting. Importantly, the ACE2-mimicking capability of these antibodies is also crucial, as it constrains viral evolution and ensures that these mAbs are not entirely evaded. Regarding mice, which lack the IGHV3-53/3-66 germline, they cannot generate a convergent response with a high amount of ACE2-mimicking NAbs, even if they are administered mRNA vaccines. It is important to note that these analyses are preliminary and intuitive, requiring further rigorous experimental validation (Extended Data Fig. 10).

## Discussion

In this study, we systematically compared the antigenicity and immunogenicity of SARS-CoV-2 XBB and JN.1 lineage from plasma to mAb resolution. Our observations reveal the robust antibody evasion and enhanced receptor ACE2 binding capabilities of the KP.3 variant, enabled by a substantial synergistic effect of F456L and Q493E mutations. Similar epistasis for ACE2 binding and evasion were also reported for “FLip” variant of XBB and other earlier variants ^12,15,49^. The ongoing evolution of JN.1 subvariants, particularly those with mutations on the A1 epitope, which are more likely to impact receptor-binding capabilities and potentially cause epistatic effects, akin to those seen in KP.3, necessitates vigilant monitoring. We underscore the importance of Class 1 or epitope A1 WT-reactive NAbs, which is also reported by a concurrent study ^50^. We additionally highlight the potential of F3 Omicron-specific NAbs elicited by Omicron reinfection cohorts in achieving broad neutralization against the JN.1 lineage.

Notably, JN.1 infection induces not only more effective NAbs, but also MBCs in both epitope groups compared to XBB. These findings support that, despite the significant cross-lineage reactivity triggered by XBB as a booster, boosters derived from JN.1 may provide superior protection against both existing and emerging JN.1 subvariants.

To enhance the generation of effective bnAbs against future antigenic drifts (such as KP.3 + A475V), it is advisable to consider developing future vaccine boosters based on KP.3 lineage, which is anticipated to become predominant in the near future due to its enhanced affinity for ACE2 and its immune evasion capabilities. For individuals who have received mRNA vaccines, the induction of WT-cross-reactive bnAbs in epitope group A1 through these boosters is particularly crucial for achieving broad-spectrum protection against both current and emerging SARS-CoV-2 variants. Additionally, if our hypothesis concerning the mechanism of heavy immune imprinting is validated, the use of a variant that demonstrates significant escape from A1 mAbs could potentially mitigate the effects of immune imprinting and effectively elicit Omicron-specific NAbs.

## Acknowledgments

We thank all volunteers who provided blood samples. This study is financially supported by the Ministry of Science and Technology of China (2023YFC3043200), Changping Laboratory (2021A0201; 2021D0102), and National Natural Science Foundation of China (32222030; 2023011477).

## Author Contributions

Y.C. designed and supervised the study. F.J. and Y.C. wrote the manuscript with inputs from all authors. A.Y., W.S., R.A., Yao W. and X.N. performed B cell sorting, single-cell VDJ sequencing experiments and data analysis. J.W. (BIOPIC), H.S., and F.J. performed and analyzed the DMS data. J.W. (Changping Laboratory) and F.S. performed the antibody expression and management. M.M. and W.W. constructed mRNA vaccines and conducted mouse immunization. Y.Y. and Youchun W. constructed the pseudotyped virus. P.W., L.Y., T.X. and W.W. performed the pseudovirus neutralization assays, ELISAs and SPR experiments. Q.G. proofread the manuscript. Y.X., X.C., Z.S. and R.J. recruited the patients.

## Data and Code Availability

Information of mAbs involved in this study is included in the supplementary tables. Other data and scripts related to this study can be downloaded from Zenedo and GitHub (to be uploaded).

## Extended Data Figure legend

**Extended Data Fig. 1.**
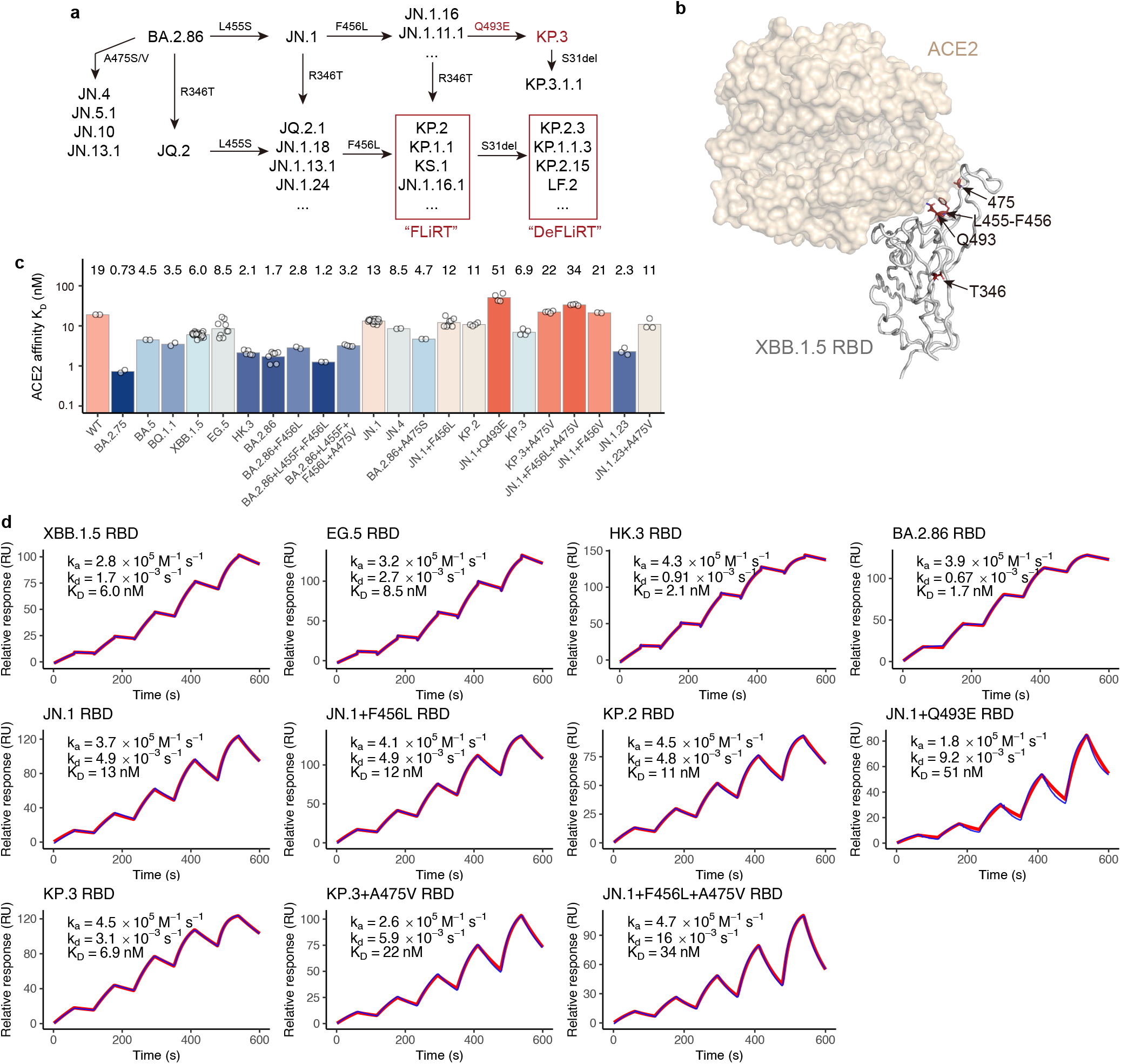
Prevalence and convergent evolution of JN.1 lineage. **a**, Schematic for the convergent evolution of BA.2.86/JN.1 lineage. **b**, Key mutated sites of BA.2.86/JN.1 lineage are indicated on the XBB.1.5 RBD structural model (PDB: 8WRL). **c**, Barplots show the affinities of additional SARS-CoV-2 variants determined by SPR. **d**, SPR sensorgrams of selected SARS-CoV-2 variants shown in Fig. 1c. Representative results of replicates are shown. Geometric mean k_a_, k_d_, and K_D_ of all replicates are labeled.

**Extended Data Fig. 2.**
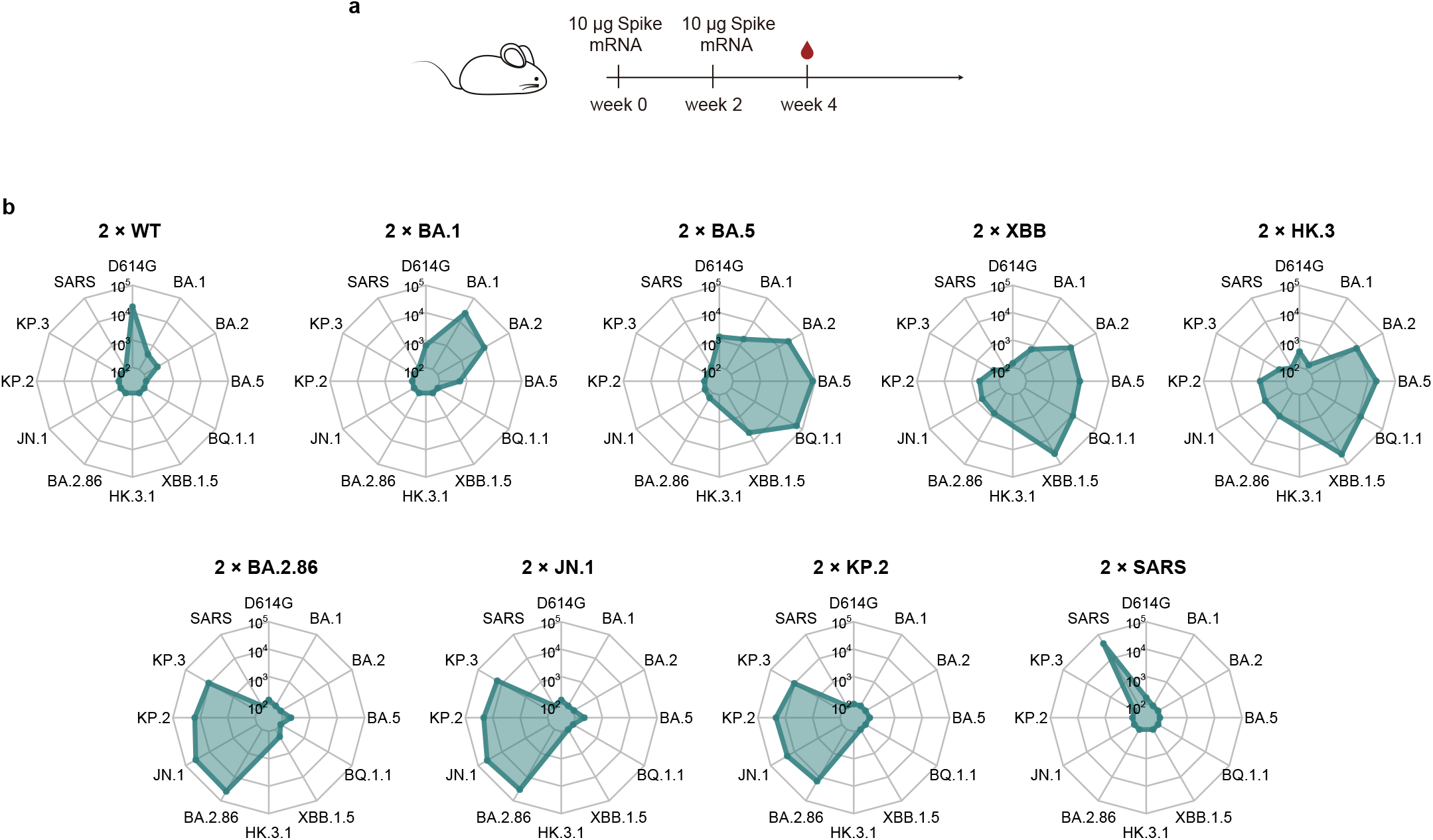
Distinct antigenicity of XBB and JN.1 in mice. **a**, Schematic for the mouse immunization experiments. **b**, Radar plots show the serum NT_50_ of mouse that received 2-dose WT, BA.1, BA.5, XBB, HK.3, BA.2.86, JN.1, KP.2, or SARS-CoV-1 Spike mRNA vaccine against eight representative SARS-CoV-2 variants.

**Extended Data Fig. 3.**
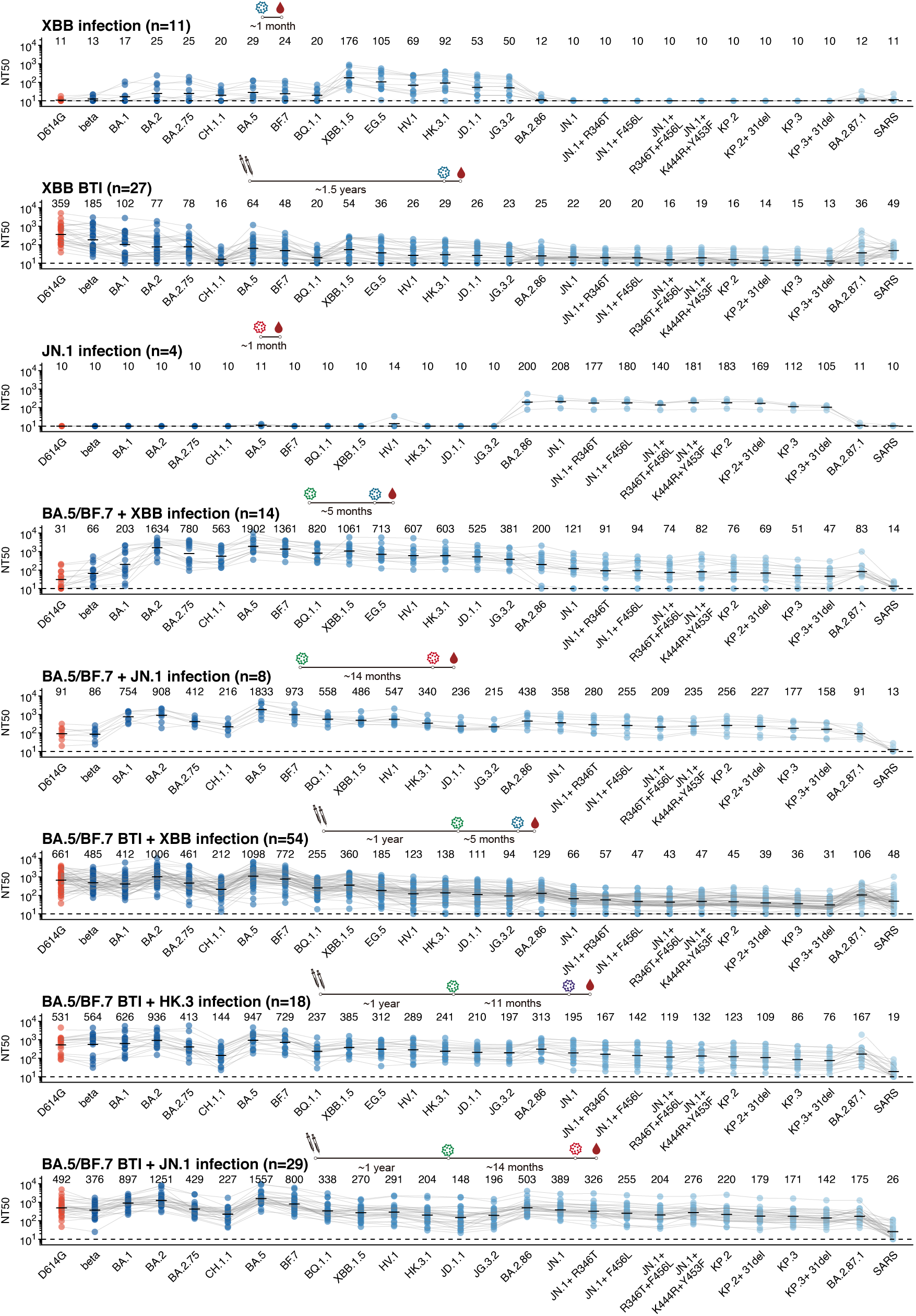
Plasma neutralization against SARS-CoV-2 variants. NT_50_ of plasma samples from all of the eight different cohorts against SARS-CoV-2 variant pseudoviruses. Plasma source cohorts and corresponding number of samples, with a schematic showing the immune history, are labeled above each panel. Dashed lines indicate the limit of detection (NT_50_ = 10). Numbers of negative samples are labeled below the dashed lines. Geometric mean titers (GMT) values are labeled as black bars and shown above each group of points. Data in Fig. 2 are displayed here again for comparison.

**Extended Data Fig. 4.**
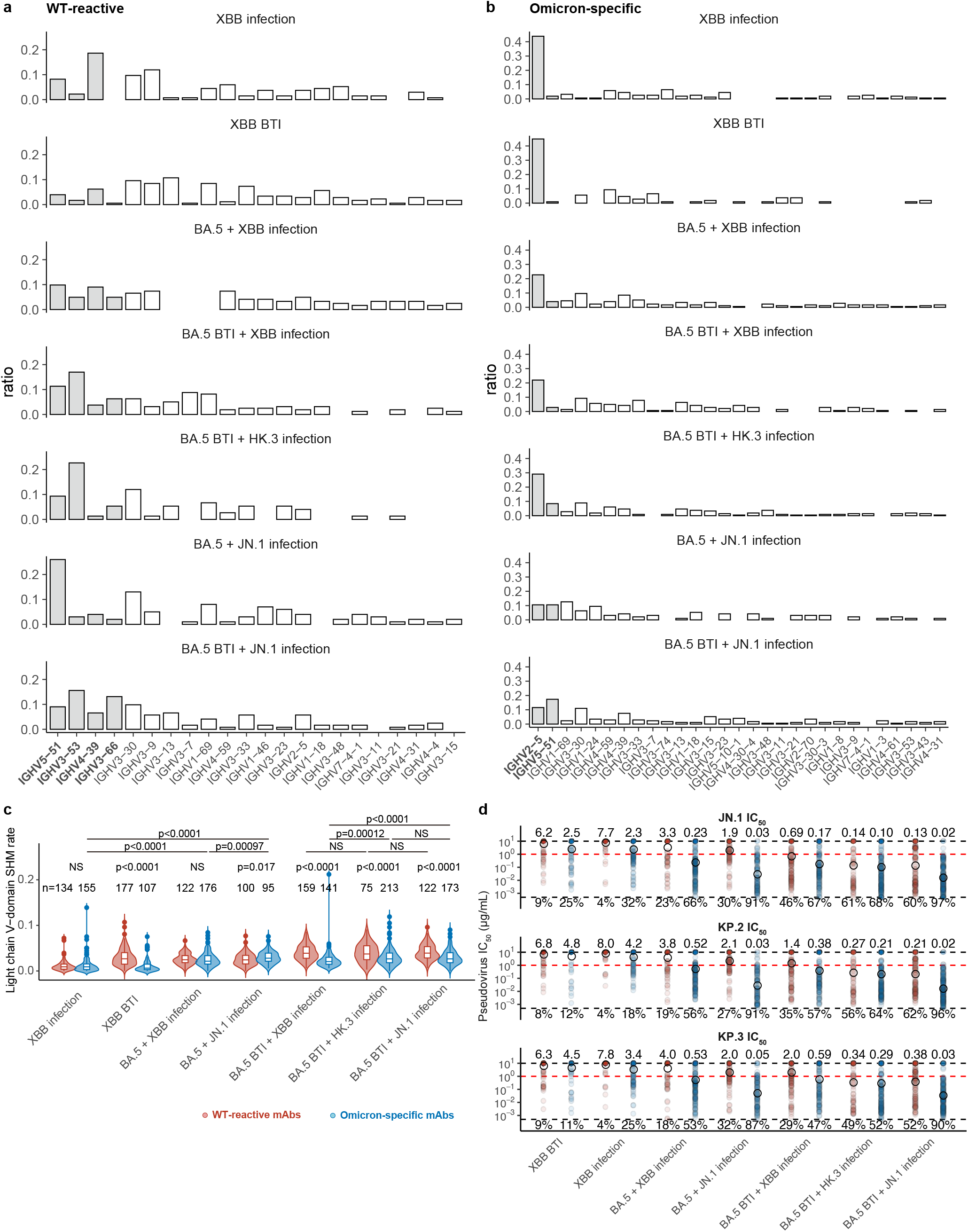
Properties of WT-reactive and Omicron-specific mAbs. **a-b**, IGHV gene distribution of WT-cross-reactive (**a**) and Omicron-specific (**b**) mAbs isolated from the seven cohorts involved in this study. **c**, Distribution of light chain SHM rate of WT-reactive and Omicron-specific antibodies isolated from different cohorts. Number of mAbs are annotated above each violin plot. Two-tailed Wilcoxon rank-sum tests are used to calculate the p-values. NS, not significant. **d**, Neutralization against JN.1, KP.2, and KP.3. Geometric mean IC_50_ values are shown as circles and annotated above the points. Black dash lines indicate limits of detection (0.005 and 10 μg/mL). Red dashed lines indicate criteria for robust neutralization (1 μg/mL). Percentage of mAbs exhibiting robust neutralization are annotated below the points.

**Extended Data Fig. 5.**
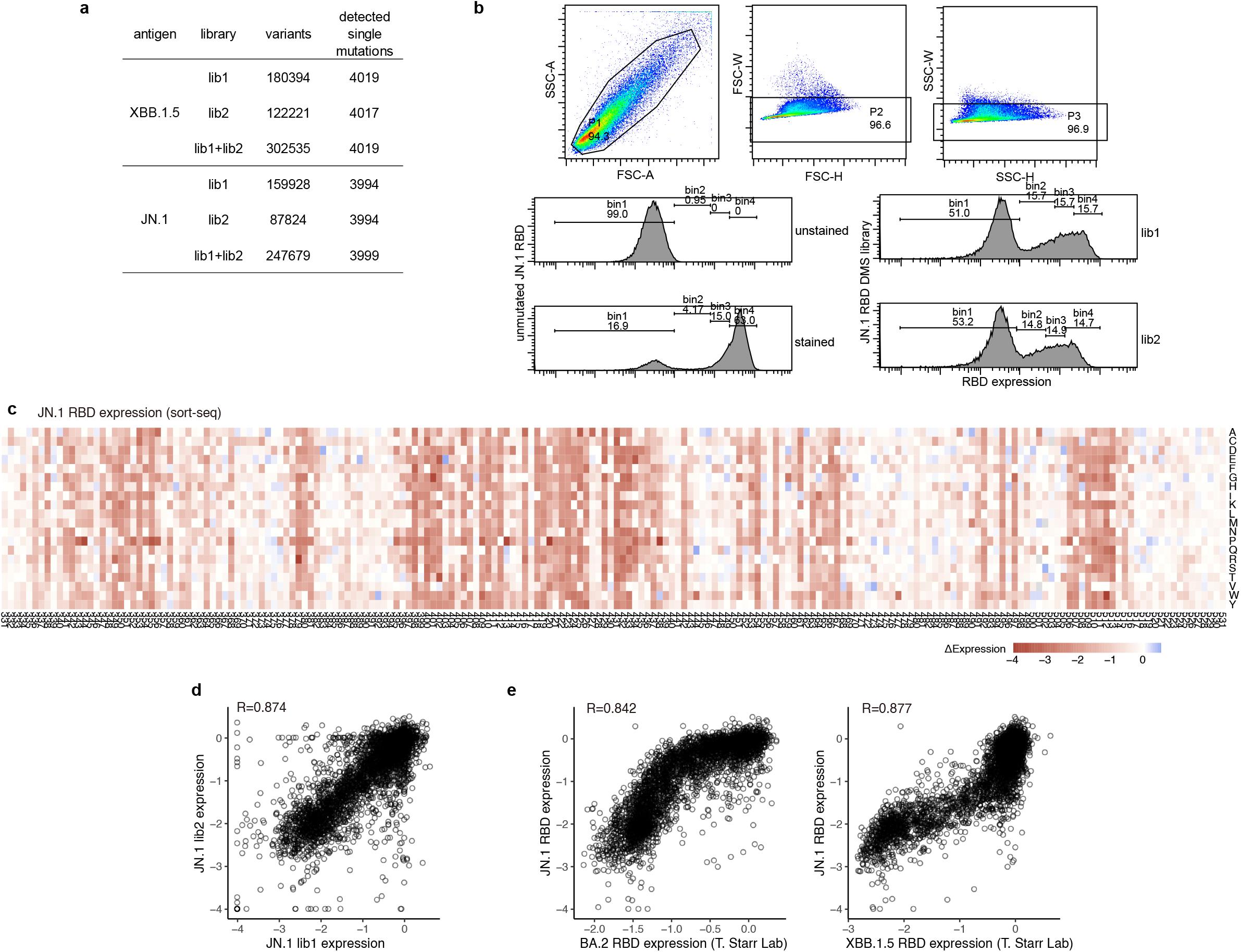
Characterization of RBD DMS mutant libraries. **a**, Number of variants and detected single mutations in the mutant libraries involved in this study. **b**, FACS diagram for Sort-seq of JN.1 mutant library to determine RBD mutant expression levels. **c**, Heatmap shows the results of DMS on RBD expression from Sort-seq. **d**, Comparison of RBD expression DMS results from two JN.1 libraries. **e**, Comparison of RBD expression DMS results between JN.1 and BA.2 (left), JN.1 and XBB.1.5 (right).

**Extended Data Fig. 6.**
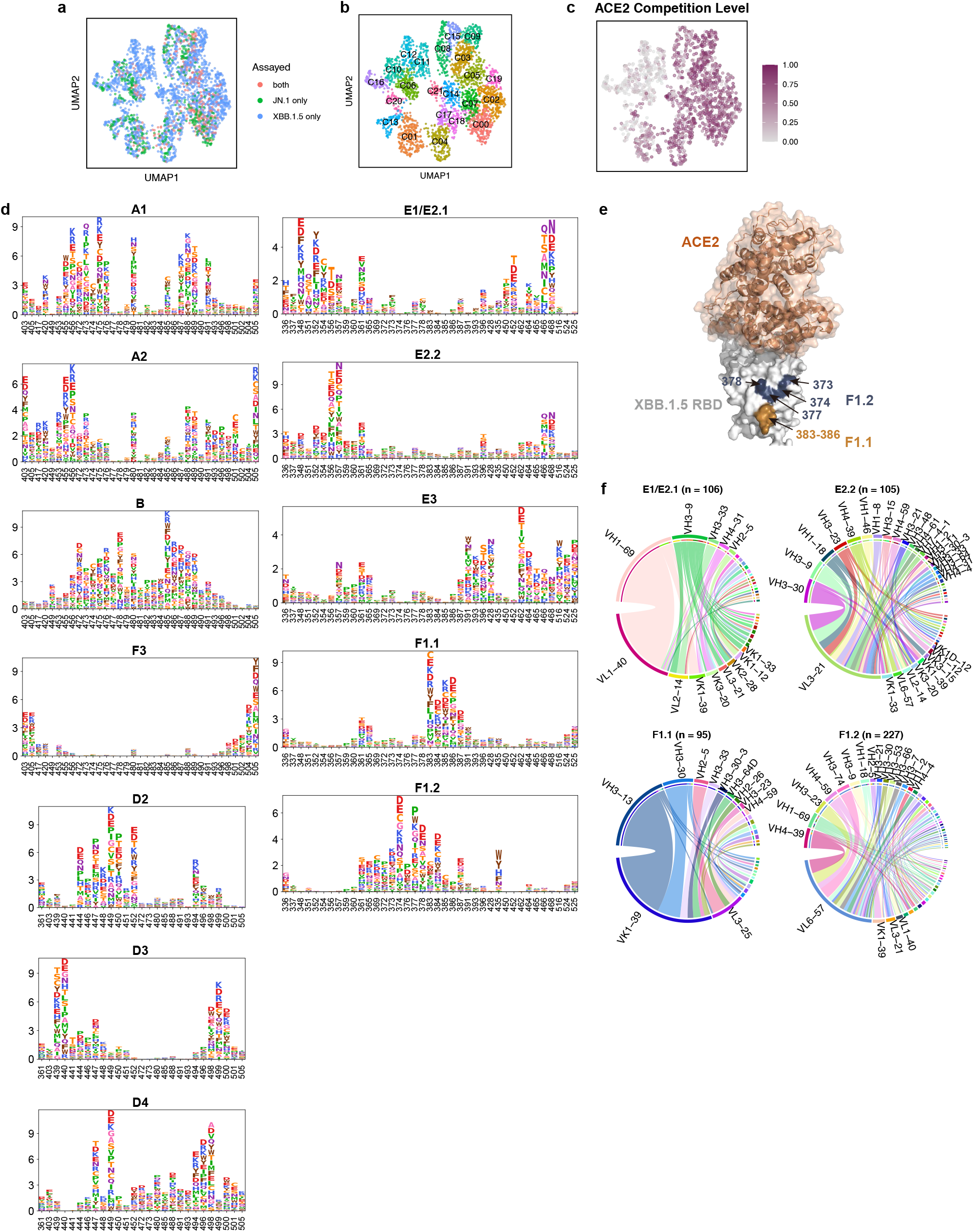
DMS-based clustering of RBD-specific mAbs. **a**, UMAP of mAbs colored by the corresponding RBD basis of DMS experiments. Some mAbs are tested in both antigen mutant libraries and the average results are used for analysis. **b**, Unsupervised clustering of DMS profiles. **c**, UMAP of mAbs colored by ACE2 competition level as determined by competition ELISA. **d**, Logo plots show average escape scores of each RBD mutation of mAbs in each epitope group. Amino acids are colored according to chemical properties. **e**, Structural model of XBB.1.5 RBD in complex of human ACE2 (PDB: 8WRL) with the key residues of epitope groups F1.1 and F1.2 highlighted. **f**, Chord diagram shows the heavy-light chain V gene pairing of mAbs isolated from in epitope groups E1/E2.1, E2.2, F1.1, and F1.2.

**Extended Data Fig. 7.**
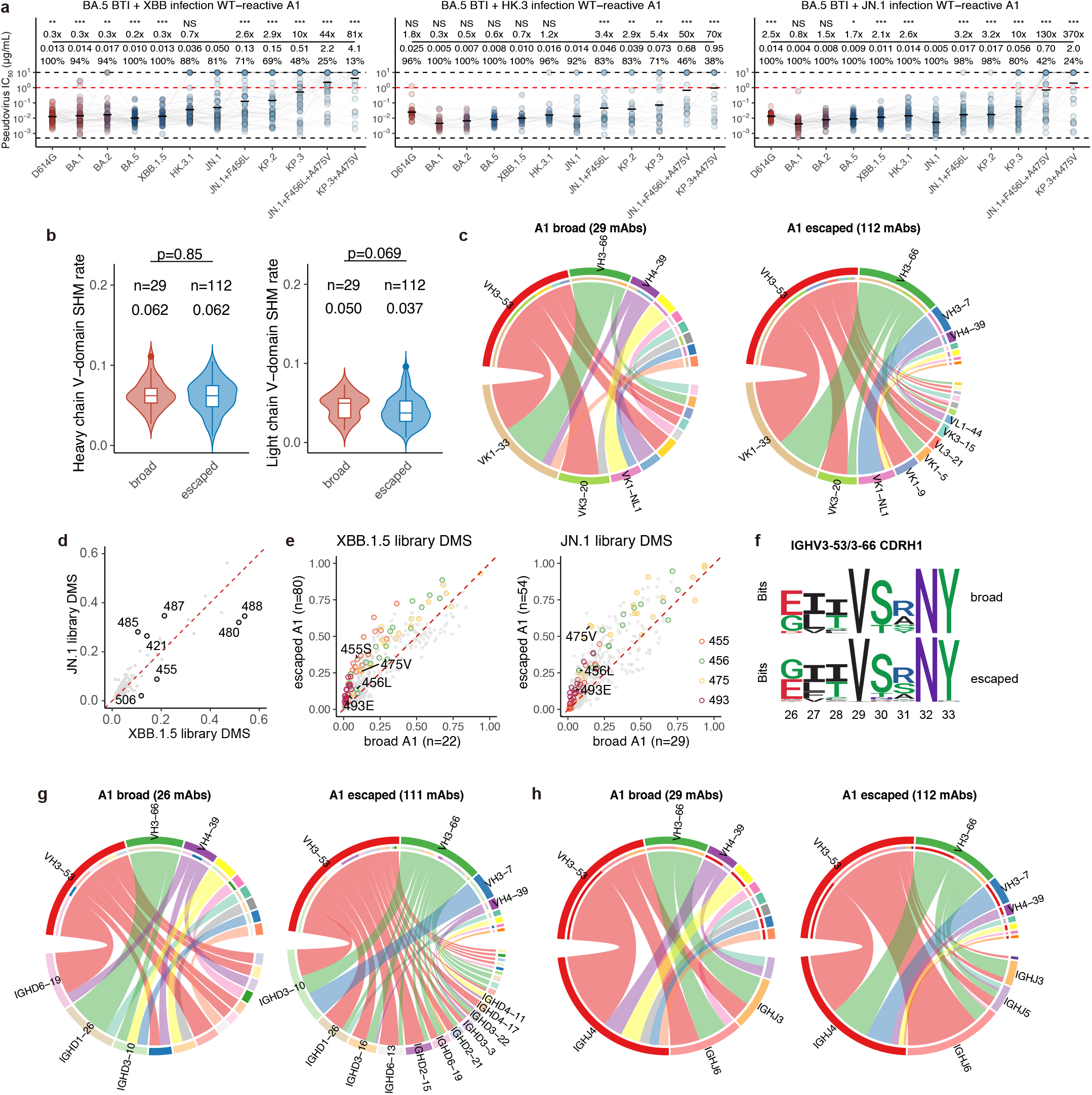
Properties of WT-reactive mAbs in epitope group A1. **a**, Neutralization of WT-reactive mAbs in epitope group A1 from three BTI + reinfection cohorts against SARS-CoV-2 variants. Geometric mean IC_50_ values are shown as circles and annotated above the points. Black dash lines indicate limits of detection (0.005 and 10 μg/mL). Red dashed lines indicate criteria for robust neutralization (1 μg/mL). Percentage of mAbs exhibiting robust neutralization, and fold-changes compared to IC_50_ against JN.1 are annotated above the points. Two-tailed Wilcoxon signed-rank tests are used to determine the p-values. *p<0.05; **p<0.01; ***p<0.001; ****p<0.0001; NS, not significant. **b**, Distribution of SHM rate of WT-reactive broadly neutralizing and escaped A1 antibodies. Number of mAbs and median SHM rates are annotated above each violin plot. Two-tailed Wilcoxon rank-sum tests are used to determine the p-values. **c**, Chord diagram shows the heavy-light chain pairing of WT-reactive broadly neutralizing and escaped A1 antibodies. **d**, Comparison of DMS site escape scores using XBB.1.5 library and JN.1 library of mAbs in epitope group A1 which were assayed in both libraries. **e**, Comparison of DMS escape scores of WT-reactive broadly neutralizing and escaped A1 antibodies. **f**, CDR-H1 motifs of IGHV3-53/3-66-encoding WT-reactive broadly neutralizing and escaped A1 antibodies. **g-h**, Chord diagram shows the heavy chain V-D (**g**) or V-J (**h**) pairing of WT-reactive broadly neutralizing and escaped A1 antibodies.

**Extended Data Fig. 8.**
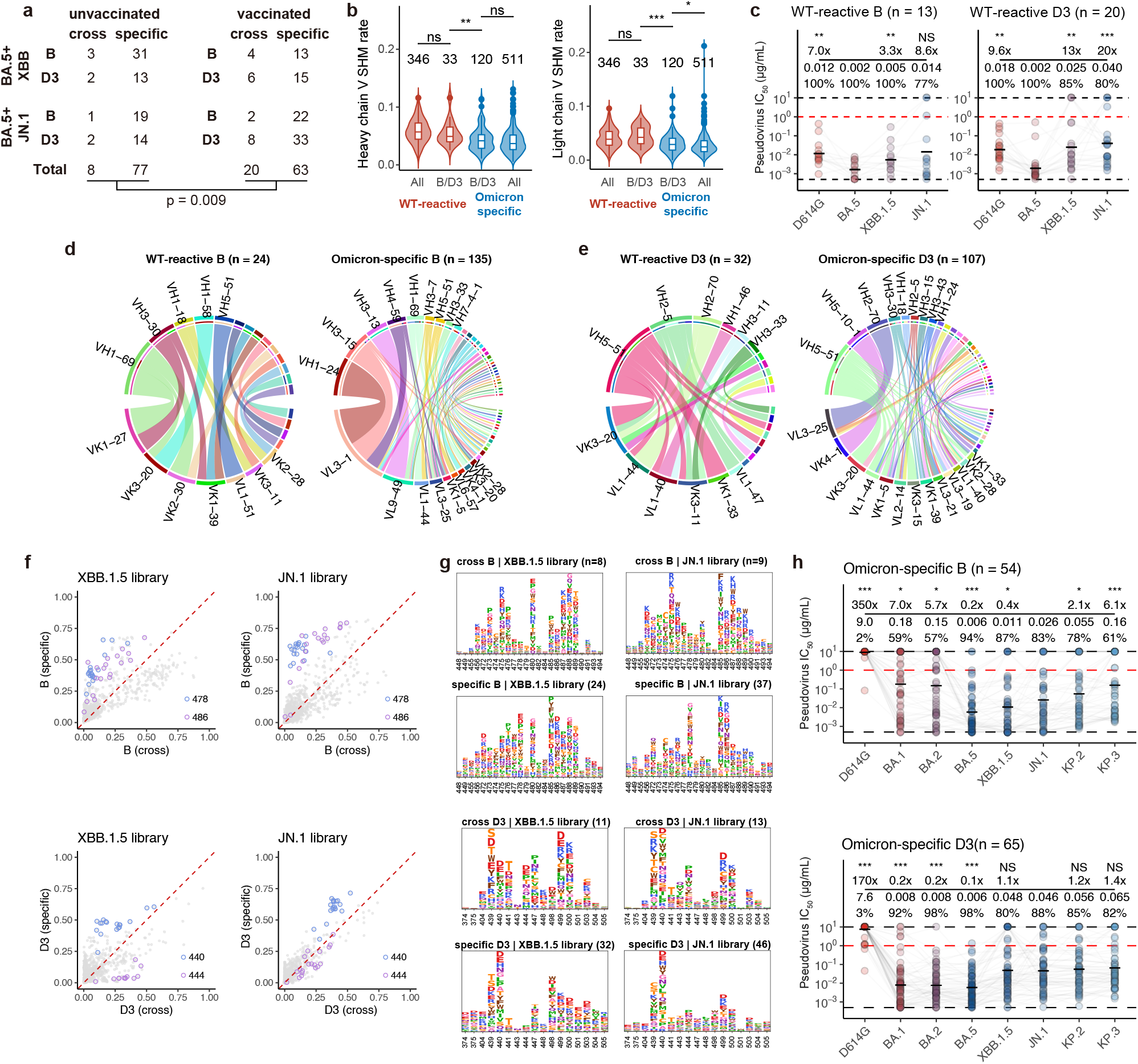
Properties of mAbs in epitope groups B and D3. **a**, Number of WT-cross-reactive and Omicron-specific mAbs in groups B and D3 from vaccinated and corresponding unvaccinated cohorts. The p-values is calculated using two-tailed hypergeometric test. **b**, Distribution of SHM rate of WT-reactive and Omicron-specific B/D3 antibodies. Number of mAbs are annotated above each violin plot. Two-tailed Wilcoxon rank-sum tests are used to determine the p-values. *p<0.05; **p<0.01; ***p<0.001; ****p<0.0001; ns, not significant. **c**, Neutralization of WT-reactive B and D3 mAbs against D614G, BA.5, XBB.1.5, and JN.1. Percentage of mAbs exhibiting robust neutralization, and fold-changes compared to IC_50_ against BA.5 are annotated above the points. **d-e**, Chord diagram shows the heavy-light chain pairing of WT-reactive and Omicron-specific B (**d**) or D3 (**e**) mAbs. **f-g**, Scatter plots (**f**) and logo plots (**g**) to compare the DMS escape scores of WT-reactive (cross) and Omicron-specific B/D3 mAbs. **h**, Neutralization of Omicron-specific B and D3 mAbs against SARS-CoV-2 variant pseudovirus. Black dash lines indicate limits of detection (0.005 and 10 μg/mL). Red dashed lines indicate criteria for robust neutralization (1 μg/mL). Percentage of mAbs exhibiting robust neutralization, and fold-changes compared to IC_50_ against JN.1 are annotated above the points. Two-tailed Wilcoxon signed-rank tests are used to determine the p-values. *p<0.05; **p<0.01; ***p<0.001; NS, not significant.

**Extended Data Fig. 9.**
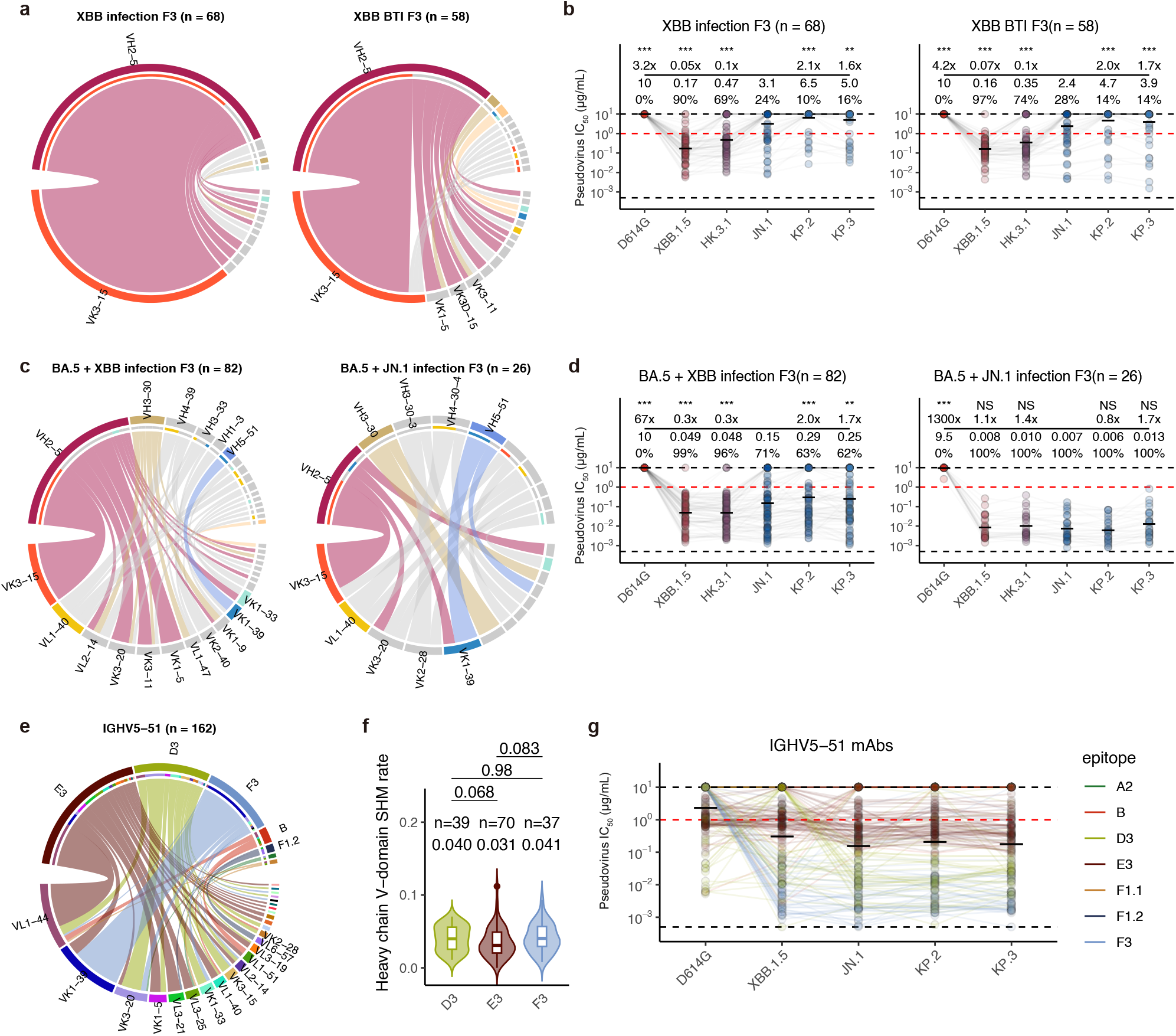
Properties of F3 and IGHV5-51 mAbs. **a**, Chord diagram shows the heavy-light chain pairing of F3 mAbs elicited by XBB infection (left) and XBB BTI (right). **b**, Neutralization of F3 mAbs s elicited by XBB infection (left) and XBB BTI (right) against SARS-CoV-2 variant pseudovirus. **c**, Chord diagram shows the heavy-light chain pairing of F3 mAbs elicited by BA.5 + XBB infection (left) and BA,5 + JN.1 infection (right). **d**, Neutralization of F3 mAbs s elicited by BA.5 + XBB infection (left) and BA,5 + JN.1 infection (right) against SARS-CoV-2 variant pseudovirus. **e**, Relationship between light chain V genes and epitope groups of IGHV5-51-encoding mAbs. **f**, Comparison of heavy chain SHM rates of IGHV5-51-encoding mAbs in epitope groups D3, E3, and F3. **g**, Neutralization of IGHV5-51-encoding mAbs in various epitope groups against D614G, XBB.1.5, JN.1, KP.2, and KP.3 pseudovirus.

**Extended Data Fig. 10.**
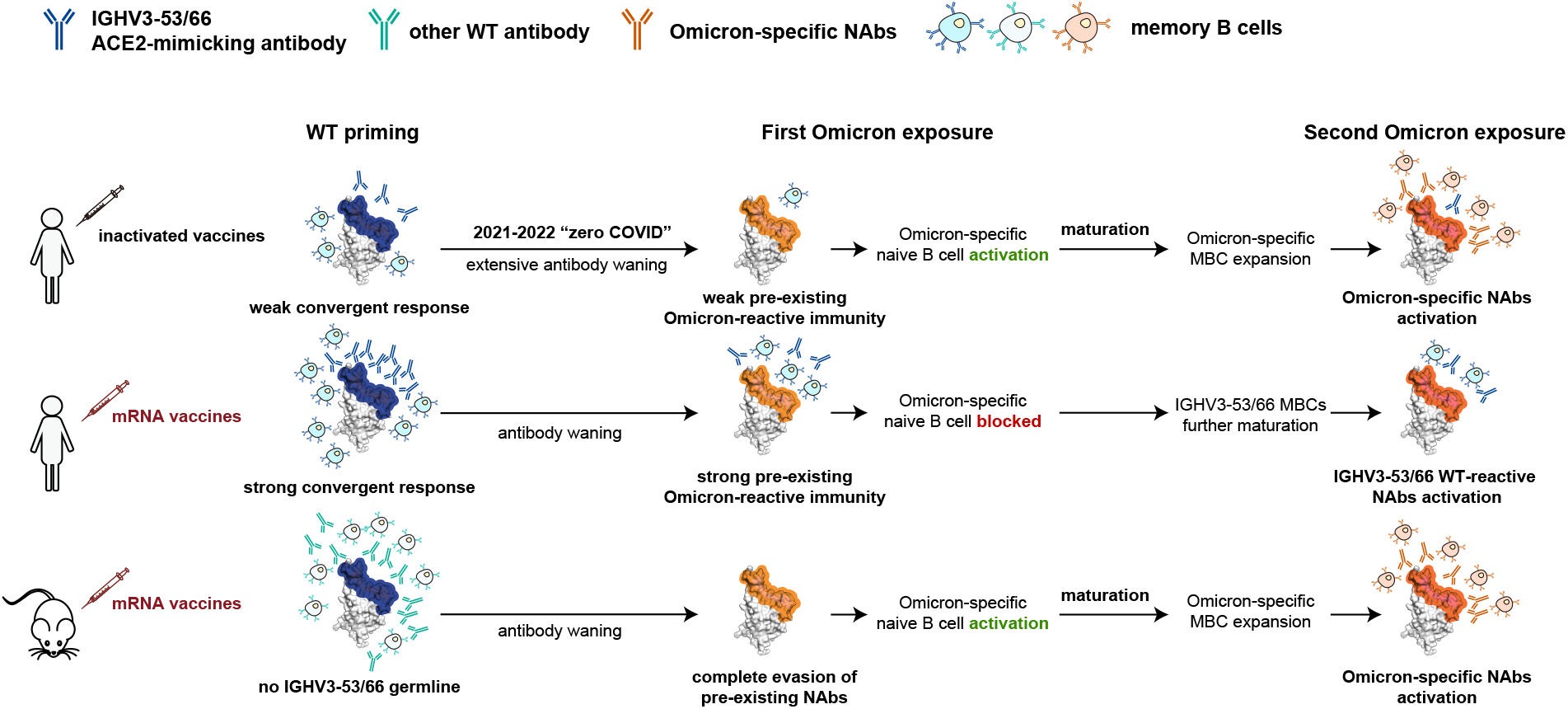
Proposed model for the distinct immune imprinting levels. Schematic for the model to explain the mRNA vaccine-induced immune imprinting.

## Supplementary Tables

**Table S1.** Information of human donors involved in this study.

**Table S2.** Information of mAbs involved in this study.

## Methods

### Plasma isolation

Blood samples were collected from individuals who had either recovered from or been re-infected with the SARS-CoV-2 Omicron BTI variant. This was conducted under the research protocol approved by the Beijing Ditan Hospital, affiliated with Capital Medical University (Ethics Committee Archiving No. LL-2021-024-02), the Tianjin Municipal Health Commission, and the Ethics Committee of Tianjin First Central Hospital (Ethics Committee Archiving No. 2022N045KY). All participants provided their agreement for the collection, storage, and use of their blood samples strictly for research purposes and the subsequent publication of related data.

Patients in the re-infection group were initially infected with the BA.5/BF.7 variants in December 2022 in Beijing and Tianjin, China ^51^. From December 1, 2022, to February 1, 2023, over 98% of the sequenced samples were identified as BA.5* (excluding BQ*), primarily consisting of the subtypes BA.5.2.48* and BF.7.14*, which were representative of the BA.5/BF.7 variants during this period. Subsequently, patients in the XBB BTI cohort and those with secondary infections in the re-infection group contracted the virus between May and June 2023. More than 90% of the sequenced samples from Beijing and Tianjin during this period corresponded to the XBB*+486P variant. These infections were confirmed using polymerase chain reaction (PCR) or antigen testing.

Whole blood was diluted in a 1:1 ratio with a solution of phosphate-buffered saline (PBS) supplemented with 2% fetal bovine serum (FBS). This was followed by Ficoll gradient centrifugation (Cytiva, 17-1440-03). After centrifugation, the plasma was collected from the upper layer, stored in aliquots at 20°C or lower, and heat-inactivated prior to subsequent experiments.

### Pseudovirus preparation and neutralization

The SARS-CoV-2 variant spike protein pseudovirus was generated using the vesicular stomatitis virus (VSV) pseudovirus packaging system as described previously ^22,52^. In addition to previously constructed variants, we additionally included “FLiRT”/KP.2 (JN.1 + R346T + F456L), KP.3 (JN.1 + F456L + Q493E), and their subvariants with S31del (SARS-CoV-2 ancestral strain numbering). The spike protein gene was codon-optimized and integrated into the pcDNA3.1 expression plasmid via the BamHI and XbaI restriction enzyme sites to augment the expression efficiency of the spike protein in mammalian cells. During pseudovirus production, the 293T cells (American Type Culture Collection (ATCC, CRL-3216)) were transfected with the SARS-CoV-2 spike protein expression plasmid. Post-transfection, these cells were infected with the G*ΔG-VSV virus (VSV-G pseudotyped virus, Kerafast) present in the cell culture supernatant. The pseudovirus was subsequently harvested and filtered from the supernatant, aliquoted, and stored at -80°C for later use.

Pseudovirus neutralization assays were performed using the Huh-7 cell line (Japan Collection of Research Bioresources [JCRB], 0403). Plasma samples were serially diluted and mixed with the pseudovirus. Following an incubation period of 1 hour at 37°C with 5% CO_2_, digested Huh-7 cells were introduced and incubated for an additional 24 hours at 37°C. The supernatant was then removed, and the mixture was incubated with D-Luciferin reagent (PerkinElmer, 6066769) in darkness for 2 minutes. The cell lysate was transferred to a detection plate, and the luminescence intensity was measured using a microplate spectrophotometer (PerkinElmer, HH3400). NT_50_ values were determined using a four-parameter logistic regression model ^53^.

### Surface plasmon resonance

SPR experiments were conducted on Biacore 8K (Cytiva) to determine the RBD-hACE2 binding affinities. Human ACE2-Fc was immobilized onto Protein A sensor chips (Cytiva). Purified SARS-CoV-2 variant RBD samples prepared in serial dilutions (6.25, 12.5, 25, 50, and 100 nM) were injected on the sensor chips. Response units were recorded by Biacore 8K Evaluation Software 3.0 (Cytiva) at room temperature. Raw response data were fitted to 1:1 binding models to determine the association and dissociation kinetic constants (k_a_ and k_d_), and binding affinities (dissociation equilibrium constant KD) using Biacore 8K Evaluation Software 3.0 (Cytiva).

In the competitive binding assays, we employed anti-His-tagged CM5 sensor chips (Cytiva) to immobilize 5 μg/mL of the RBD protein for a duration of 1 minute. Subsequently, a concentration of 20 μg/mL of antibody 1 was introduced for 2 minutes, followed by the introduction of antibody 2 at the identical concentration and for the same duration. We utilized Glycine 1.5 for the regeneration phase.

### mRNA vaccine preparation and mouse immunization

For mRNA vaccine preparation, 5′ untranslated region (UTR), target sequence, and 3′ UTR were sequentially integrated downstream of the T7 promoter within an empty PSP73 plasmid. Subsequently, a double-digestion process was employed to produce linearized DNA. This DNA served as a template for a T7 RNA polymerase-driven in vitro transcription process to generate RNA that encodes the SARS-CoV-2 S6P (F817P, A892P, A899P, A942P, K986P, V987P, R683A, and R685A) protein, according to the manufacturer’s instructions (Vazyme, DD4201). The transcriptional outputs underwent DNase I treatment for the elimination of DNA templates, followed by a purification step utilizing VAHTS RNA Clean Beads (Vazyme, N412-02). Cap 1 structure was added using Vaccinia Capping Enzyme (Vazyme, DD4109) and mRNA Cap 2′-O-methyltransferase (Vazyme, DD4110), with a subsequent purification via magnetic beads. The incorporation of Poly(A) tails was achieved with Escherichia coli Poly(A) Polymerase (Vazyme, N4111-02), culminating in another round of purification.

The mRNA was encapsulated in a functionalized lipid nanoparticle as described previously ^54^. Concisely, a solution containing ionizable lipid, DSPC, cholesterol, and PEG2000-DMG was prepared in ethanol, maintaining a molar ratio of 50:10:38.5:1.5, respectively. The mRNA was then diluted in a 50 mM citrate buffer (pH 4.0), free of RNase, to achieve a final lipid:mRNA weight ratio of 6:1. The aqueous and ethanol solutions were mixed in a 3:1 volume ratio using a microfluidic apparatus and the obtained lipid nanoparticles were then subjected to overnight dialysis. To preserve the chemical stability of the components, all samples were stored at temperatures ranging from 2 to 8 °C for up to a week. The dimensions and distribution of particle sizes of the lipid nanoparticles, as well as the encapsulation efficiency and concentration of mRNA, were meticulously assessed, revealing encapsulation rates typically between 90% and 99%.

Animal experiments were carried out under study protocols approved by Rodent Experimental Animal Management Committee of Institute of Biophysics, Chinese Academy of Sciences (SYXK2023300) and Animal Welfare Ethics Committee of HFK Biologics (HFK-AP-20210930). Female BALB/c mice, aged between six to eight weeks, were used for experiments. The mice were housed under a 12-hour light and 12-hour dark cycle, with room temperatures maintained between 20 °C and 26 °C and humidity levels maintained between 30% and 70%. mRNA vaccines were given intramuscularly at dosages of either 10 μg per mouse. Blood samples were collected 2 weeks after the final immunization, as shown in Extended Data Fig. 3a.

### Antigen-specific cell sorting and single-cell V(D)J sequencing

PBMCs and plasma were isolated from blood samples using Ficoll (Cytiva, 17-1440-03) density gradient centrifugation. B cells were enriched from PBMCs using the CD19^+^ positive selection kit (STEMCELL, 17854). The enriched B cells were then stained with RBD of the last infected variant as well as the ancestral strain RBD. B cells were also stained with antibodies against CD20 (BioLegend, 302304), CD27 (BioLegend, 302824), IgM (BioLegend, 314532), and IgD (BioLegend, 348210), and 7-AAD (Invitrogen, 00-6993-50).

B cells that were positive for last infected variant RBD (XBB.1.5, HK.3, or JN.1) and CD20, CD27, but negative for IgM, IgD and 7-AAD, were sorted. These RBD-binding B cells were subsequently subjected to single-cell V(D)J sequencing using the using the Chromium Next GEM Single Cell V(D)J Reagent Kits v1.1 according to the manufacturer’s user guide (10X Genomics, CG000208).

10X Genomics V(D)J Illumina sequencing data were assembled as BCR contigs and aligned to the GRCh38 BCR reference using Cell Ranger (v6.1.1) pipeline. For quality control, only the productive contigs and B cells with one heavy chain and one light chain were kept. The germline V(D)J genes were identified and annotated using IgBlast (v1.17.1) ^55^. SHM nucleotides and residues in the antibody variable domain were detected using Change-O toolkit (v1.2.0) ^56^.

### Expression and purification of mAbs

Antibody heavy and light chain genes were first optimized for human cell expression and synthesized by GenScript. VH and VL segments were separately inserted into plasmids (pCMV3-CH, pCMV3-CL or pCMV3-CK) through infusion (Vazyme, C112). Plasmids encoding heavy chains and light chains of antibodies were co-transfected to DH5α chemically competent cells (Tsingke, #TSC-C01-96), spread onto LB solid medium (Beyotime, #ST158) supplemented with ampicillin (Solarbio, #A1170), and single colonies cultured overnight were selected for PCR identification. Positive bacterial cultures were subjected to Sanger sequencing for verification. Finally, positive clones were selected based on sequence alignment, expanded for culture, and plasmid extraction (CWBIO #CW2105).

Expi-293F cells with a density of 0.3-0.35 × 10^6^ cells/mL were subcultured into 20 mL of culture medium (OPM Biosciences, #81075-001), sealed, and incubated at 37°C, 125 ± 5 rpm in an 8% CO_2_ atmosphere. When the cell density reached 2-3 × 10^6^ cells/mL (typically in 3 days), the cells were treated with medium to dilute the density to 2 × 10^6^ cells/mL and cultured overnight. For transfection, the antibody-encoding plasmids was diluted with 0.9% NaCl solution, mixed with polyethylenimine (PEI) transfection reagent (Yeasen, #40816ES03), and added to the cell culture. The reaction bottle was then returned to the shaker and incubated at 37°C, 8% CO_2_, and 125 ± 5 rpm. 24 hours after transfection, the matching feed solution (OPM Biosciences, #F081918-001) (1 mL/bottle) was added, and feeding was performed every other day for 6-10 days.

For antibody purification, the expression culture was centrifuged at 3000 g for 10 minutes to remove cells, and the supernatant was collected. Protein A Magnetic beads (GenScript, L00695) were added and incubated at room temperature for 2 hours, then transferred to a 24-well plate and purified using the KingFisher automated system (Thermo Fisher). The purified antibody protein was quantified using a Nanodrop (Thermo Fisher, #840-317400) and the purity confirmed by SDS-PAGE (LabLead, #P42015).

### Enzyme-linked immunosorbent assays

SARS-CoV-2 XBB.1.5, HK.3, and JN.1 RBD were individually aliquoted into a 96-well plate and incubated overnight at 4°C. The plate was then washed three times with PBST (phosphate-buffered saline with Tween-20). Subsequently, the wells were blocked with 3-5% BSA (bovine serum albumin) in PBST at 37°C for 2 hours. After another three washes with PBST, 100 μL of 1 μg/mL antibodies were added to each well and incubated for 30 minutes at room temperature. The plate was washed five times to remove unbound antibodies. Peroxidase-conjugated AffiniPure Goat Anti-Human IgG(H+L) (JACKSON, 109-035-003) was added and incubated at room temperature for 15 minutes, followed by five washes with PBST. The substrate tetramethylbenzidine (TMB) (Solarbio, 54827-17-7) was added and incubated for 10 minutes. The enzymatic reaction was halted by the addition of 2 M H_2_SO_4_. Finally, the absorbance of each well was measured at 450 nm using a microplate reader (PerkinElmer, HH3400).

### Construction of DMS libraries

Replicate DMS libraries spanning from N331 to T531 (Wuhan-Hu-1 reference numbering) of SARS-CoV-2 XBB.1.5 and JN.1 variants were constructed as outlined previously^1,2^. Initially, site-directed mutagenesis PCR with computationally designed NNS primers was conducted to generate all potential amino acid mutations on XBB.1.5 and JN.1 RBD. Then, each RBD variant was tagged with a unique 26-nucleotide (N26) barcode via PCR and assembled into Yeast surface display vector (Addgene, 166782). The XBB.1.5 and JN.1 DMS libraries were further transfected into electrocompetent DH10B cells for plasmid amplification and proceed to PacBio sequencing library preparation to decipher the association between RBD variant and corresponding N26 barcode. These enlarged DMS libraries were introduced into the EBY100 strain of *Saccharomyces cerevisiae* and screened on SD-CAA agar plates and subsequently expanded in SD-CAA liquid media, which were further preserved at -80°C after being flash-frozen in liquid nitrogen.

### Profiling of mutation effects on RBD expression

RBD expression profile for JN.1 DMS libraries was performed as previously described^2^. Briefly, yeast libraries were first recovered and propagated overnight at 30°C in SD-CAA from an original OD600 of 0.1. Then, RBD surface expression was induced by diluting the yeast cells back to SG-CAA at initial OD600 equals to 0.67 and incubating the yeasts at room temperature with mild shaking for 16 hours. Secondly, 45 OD units of induced yeasts were washed twice using PBSA (PBS supplemented with 0.2 mg/L bovine serum albumin, pH 7.4) and incubated with 1:100 diluted FITC conjugated anti-C-MYC antibody (Immunology Consultants Lab, CMYC-45F) for 1 hour at room temperature under gentle agitation. After washing with PBSA, these yeast cells were resuspended in PBSA for fluorescence-activated cell sorting (FACS). The above prepared yeasts were analyzed via BD FACSAria III cytometer by gating for single events and further partitioning into four bins according to FITC fluorescence intensity: bin 1 captured 99% of non-labelled cells while bin 2 to 4 equally divided the rest of yeasts. In total, over 25 million yeasts were collected across these four bins for each library. After sorting, yeasts from each collection tube were centrifuged for 5 minutes and resuspended in 5 mL SD-CAA. To quantify the yeast recovery rate after sorting, 10 μl of the post-sorting sample from each bin was further diluted and spread on YPD agar plates, the remaining samples were grown overnight and proceed to plasmid extraction, N26 barcode amplification and next generation sequencing.

### MACS-based antibody mutation escape profiling

High-throughput mutation escape profiling for each mAb was conducted based on magnetic-activated cell sorting (MACS) following previously reported method^1^. In brief, improperly folded RBD variants in XBB.1.5 and JN.1 DMS libraries were removed using ACE2 (Sino Biological, 10108-H08H-B) conjugated biotin binder beads (Thermo Fisher, 11533D). After washing with PBSA, the beads captured RBD expressing yeasts were released and enlarged in SD-CAA and then preserved as frozen aliquots at -80°C.

For MACS-based mutation escape profiling, the ACE2-binder yeasts were thawed in SD-CAA with shaking overnight and back-diluted into SG-CAA for RBD surface expression induction. Execution of two sequential rounds of negative selection with any given antibody eliminated specific antibody binders in libraries. Then MYC-tag-based positive selection was performed using anti-c-Myc magnetic beads (Thermo Fisher Scientific, 88843) to capture the RBD expressing yeasts in the antibody-escaping population after two rounds of negative selection.

Final obtained yeast population was washed and grown overnight in SD-CAA and submitted to plasmid extraction by 96-wells yeast plasmid extraction Kit (Coolaber, PE053). N26 barcode amplification was further conducted using obtained plasmid as the PCR template, further purified with 1X Ampure XP beads (Beckman Coulter, A63882) and subjected to single end sequencing.

### Antibody DMS data analysis

The raw sequencing data from the directed mutagenesis screening (DMS) were processed as previously described ^12,22^. Specifically, the barcode sequences detected from both the antibody-screened and reference libraries were aligned to a barcode-variant dictionary using alignparse (v0.6.2) and dms_variants (v1.4.3) tools, derived from PacBio sequencing data of the XBB.1.5 and JN.1 DMS libraries. Ambiguous barcodes were excluded during the merging of yeast libraries. Only barcodes detected more than five times in the reference library were considered for further analysis. The escape score for a variant X, present in both the screened and reference libraries, was calculated as F × (nX,ab / Nab) / (nX,ref / Nref), where F is a scaling factor to normalize the scores to a 0-1 range, and n and N represent the number of detected barcodes for variant X and the total barcodes in the antibody-screened (ab) or reference (ref) samples, respectively. For antibodies subjected to DMS with multiple replicates using different mutant libraries, the final escape score for each mutation was averaged for subsequent analyses.

We employed graph-based unsupervised clustering and embedding to assign an epitope group to each antibody and visualize them in a two-dimensional space. Initially, site escape scores (sum of mutation escape scores per residue) for each antibody were normalized to a sum of one, representing a distribution over RBD residues. The dissimilarity between two antibodies was quantified by the square root of the Jensen-Shannon divergence of the normalized escape scores. Pairwise dissimilarities for all antibodies in the dataset were computed using the SciPy module (scipy.spatial.distance.jensenshannon, v1.7.0). A k-nearest-neighbor graph was constructed using the python-igraph module (v0.9.6), and Leiden clustering was applied to assign a cluster to each antibody ^57^. Cluster names were manually annotated based on the characteristic sites in the average escape profiles of each cluster, aligning with the nomenclature of our previously published DMS dataset ^22^. To visualize the dataset in 2D, UMAP was performed based on the constructed k-nearest-neighbor graph using the umap-learn module (v0.5.2), and figures were generated using the R package ggplot2 (v3.3.3).

To compute the average immune pressure or identify escape hotspots using a collection of mAb DMS profiles, we followed a similar approach as in our previous study, incorporating four types of weights to account for the impact of each mutation on hACE2-binding affinity, RBD expression, neutralizing activity, and codon constraints at each residue, utilizing DMS data from the XBB.1.5-based DMS on ACE2 binding ^22,24^. For codon usage constraints, mutations inaccessible through single nucleotide changes were assigned a weight of zero, while others received a weight of 1.0. We used JN.1 (EPI_ISL_18373905), KP.2 (EPI_ISL_18916251), and KP.3 (EPI_ISL_19036766) to define one-nucleotide-accessible amino acid mutations. Neutralizing activity weights were calculated as -log_10_(IC_50_), with IC_50_ values below 0.0005 or above 1.0 adjusted to 0.0005 or 1.0, respectively. Raw escape scores for each antibody were normalized by the maximum score across all mutants. The weighted score for each antibody and mutation was obtained by multiplying the normalized scores by the corresponding four weights, and the final mutation-specific weighted score was the sum of scores for all antibodies in the designated set, subsequently normalized to a 0-1 range. To visualize the calculated escape maps, sequence logos were generated using the Python module logomaker (v0.8).

